# Learning from unexpected events in the neocortical microcircuit

**DOI:** 10.1101/2021.01.15.426915

**Authors:** Colleen J. Gillon, Jason E. Pina, Jérôme A. Lecoq, Ruweida Ahmed, Yazan N. Billeh, Shiella Caldejon, Peter Groblewski, Timothy M. Henley, India Kato, Eric Lee, Jennifer Luviano, Kyla Mace, Chelsea Nayan, Thuyanh V. Nguyen, Kat North, Jed Perkins, Sam Seid, Matthew T. Valley, Ali Williford, Yoshua Bengio, Timothy P. Lillicrap, Blake A. Richards, Joel Zylberberg

## Abstract

Scientists have long conjectured that the neocortex learns the structure of the environment in a predictive, hierarchical manner. According to this conjecture, expected, predictable features are differentiated from unexpected ones by comparing bottom-up and top-down streams of information. It is theorized that the neocortex then changes the representation of incoming stimuli, guided by differences in the responses to expected and unexpected events. In line with this conjecture, different responses to expected and unexpected sensory features have been observed in spiking and somatic calcium events. However, it remains unknown whether these unexpected event signals occur in the distal apical dendrites where many top-down signals are received, and whether these signals govern subsequent changes in the brain’s stimulus representations. Here, we show that both somata and distal apical dendrites of cortical pyramidal neurons exhibit distinct unexpected event signals that systematically change over days. These findings were obtained by tracking the responses of individual somata and dendritic branches of layer 2/3 and layer 5 pyramidal neurons over multiple days in primary visual cortex of awake, behaving mice using two-photon calcium imaging. Many neurons in both layers 2/3 and 5 showed large differences between their responses to expected and unexpected events. Interestingly, these responses evolved in opposite directions in the somata and distal apical dendrites. These differences between the somata and distal apical dendrites may be important for hierarchical computation, given that these two compartments tend to receive bottom-up and top-down information, respectively.

## 1 Introduction

A long-standing hypothesis in computational and systems neuroscience is that the neocortex learns a hierarchical predictive model of the world [Dayan et al., 1995; Friston and Kiebel, 2009; Hawkins and Blakeslee, 2004; Larochelle and Hinton, 2010; Press et al., 2020; Rao and Ballard, 1999; Spratling, 2017; Whittington and Bogacz, 2017]. This hypothesis postulates that learned top-down predictions (i.e., signals from associative regions to sensory regions) are compared to bottom-up signals (i.e., signals from sensory regions to associative regions) (Fig. 1A). Unexpected stimulus events should then induce differences between these signals and, in turn, drive learning. In these models, learning occurs at all stages of the hierarchy, and not just at the earliest or latest stages. Theoretical support for this hypothesis comes from computational studies showing that hierarchical models that learn by comparing top-down signals to bottom-up signals enable artificial neural networks (ANNs) to learn useful representations that capture the statistical structure of the data on which they are trained [Chen et al., 2020; Devlin et al., 2018; Grill et al., 2020; Lotter et al., 2016; van den Oord et al., 2018; Wayne et al., 2018]. Moreover, ANNs trained in this manner reproduce the representations observed in the neocortex better than ANNs trained purely by supervised learning based on categorical labels [Bakhtiari et al., 2021; Christensen and Zylberberg, 2020; Higgins et al., 2017].

**Figure 1:**
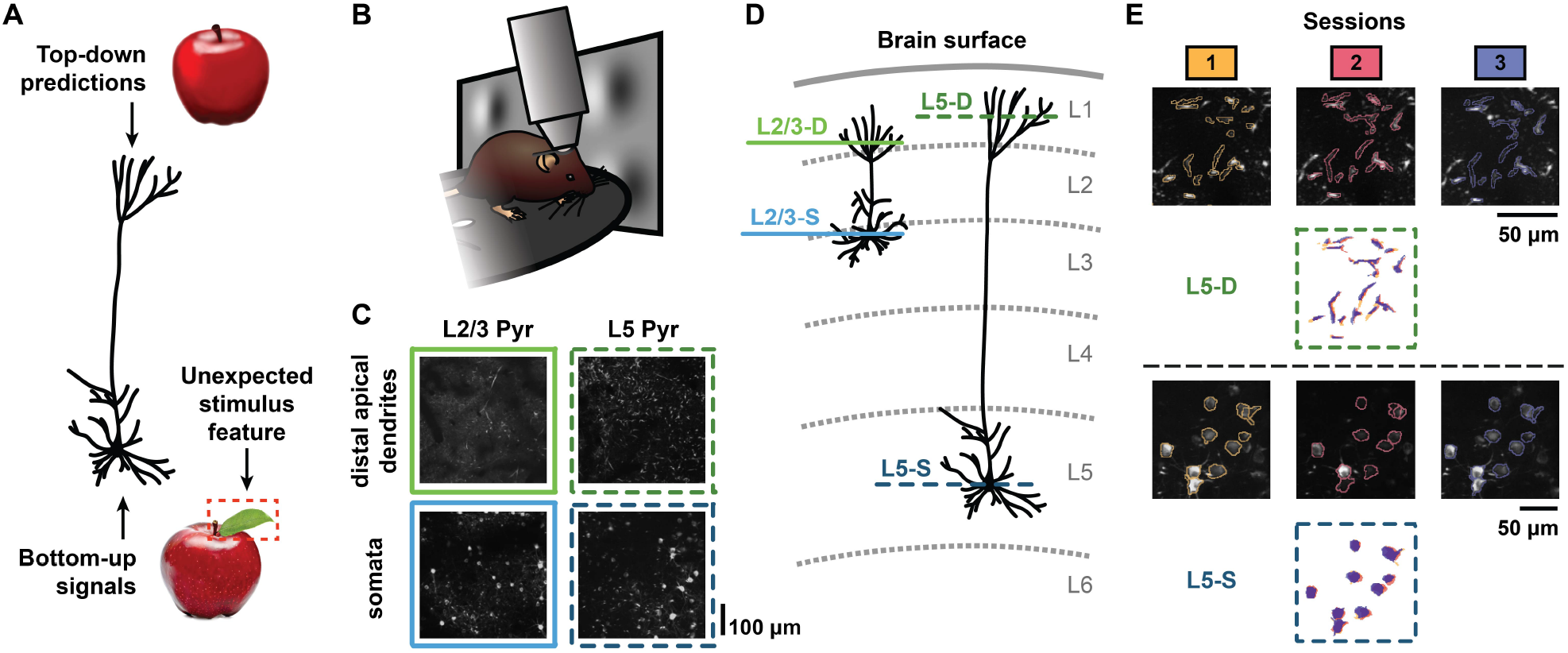
Illustration of experimental methods. **(A)** Schematic illustration of how a hierarchical predictive model might be implemented through the morphology of pyramidal neurons. The neuron receives top-down predictions (e.g., corresponding to an “expected” model of an apple, *top*) at the distal apical dendrites that are compared to bottom-up stimulus information received perisomatically (e.g., corresponding to an actual image of an apple, *bottom*). In this cartoon example, the incoming stimulus contains an unexpected feature (a leaf) not captured by the top-down predictive model. **(B)** Experimental setup schematic. Awake, behaving mice were head-fixed under a two-photon microscope objective while passively viewing the stimuli. The mice were able to run freely on a rotating disc. **(C)** Example maximum-projection images from two-photon recordings for each of the four imaging planes: layer 2/3 distal apical dendrites (L2/3-D), layer 5 distal apical dendrites (L5-D), layer 2/3 somata (L2/3-S), and layer 5 somata (L5-S) (2–3 mice per plane, *n* = 11 mice in total; see Materials & Methods). The corresponding recordings are shown in Supp. Video 1. **(D)** Schematic illustration displaying the four imaging planes from (C) within the cortical column. The coloring and style schemes of the horizontal lines depicting the imaging planes here are used throughout all of the figures. **(E)** Tracked region of interest (ROI) examples for both L5-D (*top*) and L5-S (*bottom*). Maximum-projection images for each imaging session (1, 2, or 3, as indicated), each performed on a different day, are overlaid with contours of the matched segmented ROI masks. Below the images, the matched segmented ROI masks for all three sessions are superimposed. See also Fig. S1.

What would be the observable signatures of this type of hierarchical predictive learning, in which learning is guided by unexpected sensory events? There are at least three signatures that one might expect: (1) There should be distinct responses to expected and unexpected stimuli. If the brain does not distinguish between expected and unexpected events, there is no way to specifically learn from the *unexpected* events. (2) As the circuit learns about stimuli, the responses to both expected and unexpected stimuli should change in a long-lasting manner. These changes in stimulus responses are a necessary consequence of learning modifying the stimulus representations. (3) There should be differences between the manner in which top-down and bottom-up driven responses change during learning. This follows from the idea that a hierarchical model is being learned, since hierarchy implies a distinct role for top-down and bottom-up information.

Previous work has provided partial evidence for these observable signatures of hierarchical predictive learning. First, there is a very large body of work showing distinct responses to expected and unexpected stimuli in multiple species and brain regions [Fiser et al., 2016; Garrido et al., 2009; Keller et al., 2012; Kumaran and Maguire, 2006; Orlova et al., 2020; Zmarz and Keller, 2016], thus supporting the first observable signature. However, there are still significant unknowns: e.g., do such responses evolve differently in different compartments of neurons? Second, there is some research suggesting that responses to unexpected stimuli change with exposure [Homann et al., 2017], supporting the second observable signature. Yet, this has only been shown over short time scales, such as a single experimental session. Third, there are a few studies showing that top-down projections carry distinct information to sensory areas [Fiser et al., 2016; Jordan and Keller, 2020; Orlova et al., 2020], partially supporting the third observable signature. Nonetheless, it remains unknown whether changes in neural responses driven by top-down versus bottom-up signals show distinct changes over learning. Thus, the goal of this paper is to fill these gaps by concretely looking for all three of these signatures together in a systematic study.

Here, we performed chronic two-photon calcium imaging of layer 2/3 and layer 5 pyramidal neurons at both the cell bodies and the distal apical dendrites in the primary visual cortex of awake, behaving mice over multiple days (Fig. 1B–D; Supp. Videos 1-3). These imaging planes were chosen since top-down signals largely impinge on the distal apical dendrites within cortical layer 1, while bottom-up signals largely impinge on the perisomatic compartments in deeper layers [Budd, 1998; Larkum, 2013a,b]. During the recordings, the animals were exposed to randomly oriented visual stimuli with both expected and unexpected statistical properties (Supp. Videos 4–5). Altogether, this approach allowed us to track the responses of both individual cell bodies and individual distal apical dendritic branches over multiple days (Fig. 1E), during which the animals were provided with more exposure to unexpected events. The resulting data showed evidence corroborating all three of the signatures of hierarchical predictive learning above, supporting the hypothesis that the visual cortex learns from unexpected events using a hierarchical model. Moreover, we observed interesting differences between the distal apical dendrites and somata. Whereas somatic compartments showed a decrease in differential sensitivity to expected versus unexpected visual stimuli over days, distal apical dendrites showed an increase in differential sensitivity. This suggests that there may be important differences in the functional roles of the somatic and distal apical compartments in hierarchical predictive learning in the neocortex.

## 2 Results

### 2.1 Imaging dendrite segments and cell bodies over multiple days

To monitor the integration of top-down and bottom-up signals by supra- and sub-granular pyramidal neurons over multiple days, we performed two-photon calcium imaging in Cux2-CreERT2 mice or Rbp4-Cre KL100 mice that expressed GCaMP6f in layer 2/3 or layer 5 pyramidal neurons, respectively. We performed this imaging either at layer 1 of cortex (50–75 *µ*m depth for layer 2/3 and 20 *µ*m depth for layer 5), thereby observing the distal apical dendrites, or at the layer in which the cell bodies were located (175 *µ*m depth for layer 2/3 and 375 *µ*m depth for layer 5) (Fig. 1C–D; Supp. Video 1). This gave us four different imaging conditions: layer 2/3 distal apical dendrites (L2/3-D), layer 2/3 somata (L2/3-S), layer 5 distal apical dendrites (L5-D), and layer 5 somata (L5-S). GCaMP6f fluorescence tracks calcium influx into cells, but it should be noted that the cause of calcium influx in the somatic and distal apical compartments may be different, with somatic signals largely reflecting closely-spaced groups of action potentials [Huang et al., 2021] and dendritic signals largely reflecting non-linear dendritic events like plateau potentials [Murayama et al., 2009]. Thus, in both cases we are tracking a proxy for neural activity, but it is important to be aware that the underlying physiological cause of the signal likely differs between the two compartments.

Imaging was performed in primary visual cortex (VisP). During the experiments, the animal’s head was fixed in place under the microscope objective, ensuring the stability of our recordings. We extracted regions of interest (ROIs) in each imaging plane [de Vries et al., 2020; Inan et al., 2017, 2021], corresponding to individual distal apical dendrite segments or to individual cell bodies, depending on the imaging plane. Each animal went through three imaging sessions, each performed on a different day, and we used a matching algorithm to identify the same ROIs across sessions (Fig. 1E, S1).

Thanks to a very conservative quality control pipeline (see Materials & Methods), signal-to-noise ratio (SNR), Δ*F/F* magnitudes, and number of ROIs were stable over all three sessions in both layer 2/3 and layer 5 cell bodies and dendrites (Fig. S2). Importantly, the ROI extraction algorithm for the dendritic recordings enabled the identification of spatially discontinuous ROIs [Inan et al., 2017, 2021], reducing the risk that single dendritic compartments were split into multiple ROIs. This is supported by the observation that in both the somatic and the dendritic compartments, very few pairs of ROIs showed very high correlations in their responses (Fig. S2D). Moreover, while differences in background fluorescence levels were observable between imaging planes (Fig. 1C), these did not confound our analyses for two reasons. First, we only compared Δ*F/F* levels over days within each imaging plane, not between imaging planes. Second, our analysis pipeline estimated Δ*F/F* using a rolling baseline, so that changes in overall fluorescence would not impact our analyses (see Materials & Methods).

During these imaging sessions, we tracked the mouse’s movements on a running disc (Supp. Video 2) as well as its pupil diameter with an infrared camera (Supp. Video 3). We obtained calcium imaging data for 11 mice (L2/3-D: *n* = 2, L2/3-S: *n* = 3, L5-D: *n* = 3, L5-S: *n* = 3). The full dataset is freely available online in the DANDI Archive (see Materials & Methods).

### 2.2 Cortical neurons respond differently to expected and unexpected stimuli

To explore the responses of cortical neurons to expected and unexpected sensory events, we designed a sequential visual stimulus inspired by previous work [Homann et al., 2017]. This stimulus had a predictable global structure, but stochastic local properties. Thanks to the predictable global structure we could randomly insert “unexpected” events, i.e., stimulus events that violated the predictable global pattern. Mice were exposed to this stimulus over multiple sessions, each occurring on different days, enabling us to observe changes in their neurons’ responses to expected and unexpected sensory events.

To build a predictable global structure with some local stochasticity, we used image frames composed of randomly placed Gabor patches, assembled into five-frame sequences (*A-B-C-D-G*). Other than *G*, which was uniformly gray, each frame was defined by the locations of its Gabor patches: e.g., the locations of the Gabor patches were the same for all *A* frames for a given session, but differed between *A* and *B* frames. These Gabor patch locations were redrawn for each session, and sampled uniformly over the visual field. As a result, the locations were different in each session. Additionally, within each repeat of the sequence (*A-B-C-D-G*), the orientations of each of the Gabor patches were drawn randomly from the same distribution centered around the same mean orientation (Fig. 2A, Supp. Video 4), but the mean orientation varied from sequence to sequence. This meant that the luminance patterns at each spatial location were different for each repeat of the *A-B-C-D-G* sequence. However, because all sequences shared a global pattern wherein orientations were drawn from the same distribution across frames, knowing the orientations of the Gabors from one frame of the sequence would enable clear predictions about the orientations of the Gabors in the subsequent frames. Importantly, given these stimulus design features, the same set of images was never repeated. This reduced the risk of accommodation effects, which could cause changes in neuronal responses via mechanisms other than learning. Nonetheless, the sequences had predictable global properties that would allow an observer to form expectations about upcoming frames. Thus, the animals could learn the “rules” underlying the stimuli with increasing exposure and thereby form expectations for what should happen next. It is important to note that we cannot say with certainty whether the animals actually expected the stimulus sequences. We can, however, say that they were provided with substantial experience with which to form such expectations. For that reason, we call these *A-B-C-D-G* sequences “expected”. See Materials & Methods for a more detailed description of the stimulus properties.

**Figure 2:**
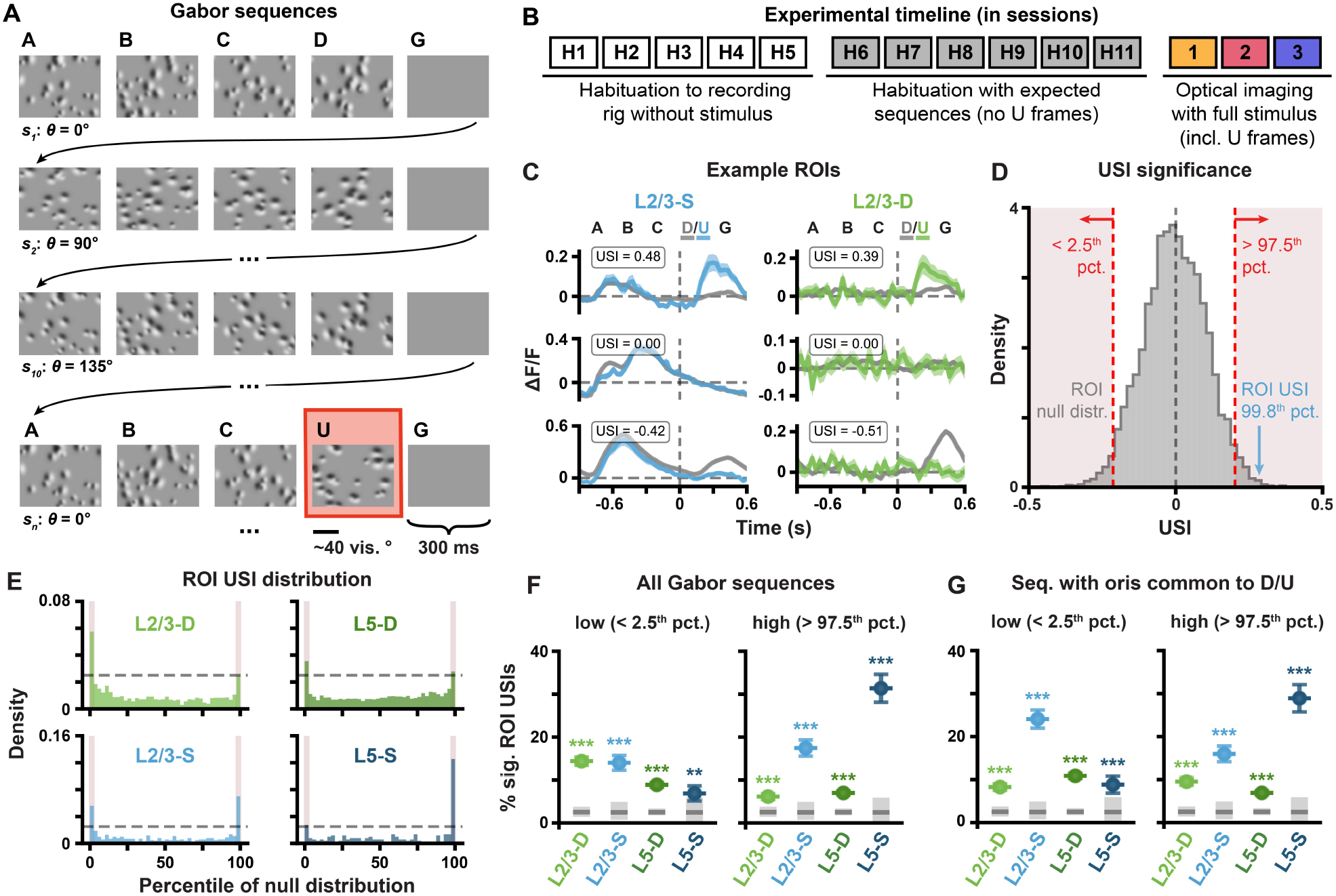
Δ*F/F* responses to unexpected stimuli reflected in the first imaging session. **(A)** Example Gabor sequences. Each frame lasted 300 ms. The mean orientation *θ* of the Gabor patches in each sequence *s_i_* was randomly chosen from {0^°^, 45^°^, 90^°^, 135^◦^}. An unexpected *U* frame, with a mean orientation rotated by 90^°^ with respect to the other frames in the sequence, is highlighted in red. See Sec. 2.2, Materials & Methods and Supp. Video 4 for more details. **(B)** Experimental timeline, showing both habituation and imaging sessions. Note that each session occurred on a different day. Optical imaging of neuronal activity was not performed during *H1–H11*. **(C)** Example session 1 Δ*F/F* response traces for individual L2/3-S (*left*) and L2/3-D (*right*) ROIs with high (*top*), null (*middle*) or low (*bottom*) USIs. Mean *±* standard error of the mean (SEM) Δ*F/F* across Gabor sequences is plotted. Dashed vertical lines mark onset of D/U frames. **(D)** Example USI null distribution for one ROI from L2/3-S in session 1, generated by shuffling *D-G* and *U-G* labels for the same ROI and recomputing the shuffled USIs 10^4^ times. Significant regions highlighted in red, and true USI value labelled in blue. **(E)** USI percentile distributions for each plane for all session 1 ROIs. Dashed horizontal lines depict null hypotheses (i.e., uniform distribution). Significant percentiles are marked with red highlights (*p <* 0.05, shuffle test as shown in (D)). **(F)** Percentage *±* bootstrapped standard deviation (SD) of significant USIs for all segmented ROIs in session 1 for each plane. All sequences (any mean orientation) are included in the analysis. **(G)** Same as (F), but restricted to the Gabor sequences with mean orientations shared between *D* and *U* frames {90^°^, 135^◦^}. *: *p <* 0.05, **: *p <* 0.01, ***: *p <* 0.001 (two-tailed, corrected). See Table S1 for details of statistical tests and precise p-values for all comparisons.

To help the animals form such expectations, before the first calcium imaging session, the mice were habituated to *A-B-C-D-G* sequences over multiple sessions, each on a different day, without any violations of the predictable structure (Fig. 2B). After habituation, and during calcium imaging, the stimuli were broken up into approximately 30 blocks of randomly determined durations, each composed of repeated *A-B-C-D-G* sequences, as before. However, instead of comprising only expected sequences, each block ended with “unexpected” *A-B-C-U-G* sequences. In these sequences, the fourth frame, *D*, was replaced with an unexpected *U* frame, which had different Gabor locations and orientations. Specifically, the newly introduced *U* frames had unique random locations and the orientations of the Gabor patches were resampled and shifted by 90*^◦^* on average with respect to the preceding *A-B-C* frames. As such, the *U* frames strongly violated potential expectations about both Gabor patch locations and orientations. These “unexpected” sequences comprised approximately 7% of the sequences presented to the mice during the imaging sessions.

ROIs in VisP exhibited clear responses to both the onset of the sequences and the final Gabor frames (Fig. S3), so we compared the responses of the ROIs to the unexpected *U* frames and the expected *D* frames. First, we examined the average Δ*F/F* signals in each of the four imaging conditions: L2/3-D, L2/3-S, L5-D, L5-S. We observed that some ROIs had clearly different responses to the expected and unexpected frames (Fig. 2C, *top* and *bottom* traces). To quantify the difference in responses to the expected versus the unexpected frames, we calculated an “***u***nexpected event ***s***electivity ***i***ndex” (USI) by subtracting the mean responses to the expected from the unexpected stimulus events, and scaling this value by a factor of their variances (Equation 1). We then examined the USIs to see whether they indicated that the circuit treated the expected and unexpected frames differently. We found that many more ROIs than would arise by chance had negative or strongly positive USIs, as has been previously observed [Keller et al., 2012]. To determine chance levels, we constructed null distributions non-parametrically for each ROI by shuffling the “expected” and “unexpected” labels for the stimulus frames 10^4^ times, each time recomputing the USI on the shuffled data (Fig. 2D; see Materials & Methods). These shuffles yielded a null distribution over USI values for each ROI that reflected the null hypothesis, according to which there was no difference in an ROI’s responses to expected and unexpected events. We then identified the percentile of each ROI’s real USI within its own null distribution: ROI USIs below the 2.5th percentile or above the 97.5th percentile were labelled as statistically significant (Fig. 2D). Across the population of ROIs, in both L2/3 and L5 somata and dendrites, there were far more significant USIs than would be predicted by chance (Fig. 2E–F). This effect was consistent across individual mice, with 10 of the 11 animals showing a statistically significant effect (Fig. S4A). Notably, when we restricted this analysis to sequences whose mean Gabor patch orientations occurred for both *D* and *U* frames, namely 90*^◦^* and 135*^◦^*, the USI percentages remained largely the same, meaning that USI patterns did not reflect ROI preferences for specific orientations of the Gabor patches (Fig. 2G). Thus, the response differences we observed were unlikely to be a result of the differences in the orientations of the Gabor patches in the *D* and *U* frames. Together, these data indicate that the neurons and dendrites in primary visual cortex respond systematically differently to expected and unexpected frames, in line with the first observable signature of predictive learning discussed above.

We next wondered whether the differences in the responses to expected and unexpected frames could have been driven by differences in the animals’ behavior. There is a growing body of evidence showing that responses in mouse visual cortex are affected by behaviors like running and pupil dilation [Niell and Stryker, 2010; Salkoff et al., 2020; Stringer et al., 2019]. Therefore, it was important to ask whether the mice altered their behavior in response to the unexpected stimulus frames. If so, these behavioral differences could be reflected in the neuronal responses in visual cortex, confounding our interpretation that the differences in neuronal response were due to the expected versus unexpected nature of the stimulus. To test this possibility, we compared the animals’ running velocities and pupil dilation during the expected *D* frames and the unexpected *U* frames (Fig. 3A). We found no difference in either running velocity or pupil dilation for *D* versus *U* frames (Fig. 3B), suggesting that behavioral changes are not a major confound in our analyses. Altogether, these data confirm the first observable signature of hierarchical predictive learning introduced above, i.e., that expected and unexpected stimuli are represented differently within the neocortical microcircuit.

**Figure 3:**
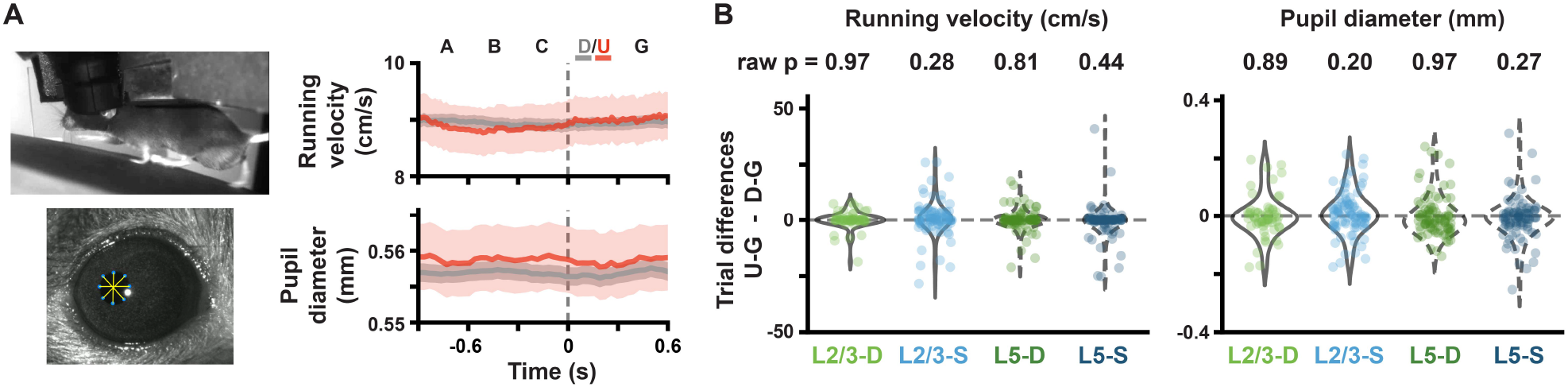
Absence of behavioral responses to unexpected stimuli. **(A)** (*Left*) Example frames of a mouse running (*top*), and of a mouse pupil with tracking markers (*bottom*). (*Right*) Running velocity and pupil diameter traces aggregated across mice (mean *±* SEM across Gabor sequences) for expected (gray) and unexpected (red) sequences. Note that the smaller SEM is due to the greater number of expected sequences, compared to unexpected ones. Dashed vertical lines mark onset of D/U frames. **(B)** Block-by-block running velocity (*left*) and pupil diameter (*right*) differences between unexpected (*U-G*) and expected (*D-G*) frames. Raw two-tailed p-values (not corrected for multiple comparisons) are shown. *: *p <* 0.05, **: *p <* 0.01, ***: *p <* 0.001 (two-tailed, corrected). See Table S1 for details of statistical tests and precise p-values for all comparisons.

### 2.3 Responses to expected and unexpected stimuli evolve over days and differ between the somata and distal apical dendrites

To probe learning, we compared the neural responses to expected and unexpected stimuli over three sessions spread across multiple days. Importantly, unsupervised learning—wherein a system learns about stimuli merely through exposure to them [Beaulieu and Cynader, 1990; Lotter et al., 2016; van den Oord et al., 2018; Woloszyn and Sheinberg, 2012; Zylberberg et al., 2011]—is not necessarily associated with any behavioral changes. As such, experimentally observing unsupervised learning requires observing changes in neural representations as animals gain experience with sensory stimuli. Therefore, we examined the evolution of the neuronal responses to expected and unexpected stimuli over the three different days of calcium imaging. This analysis made use of our ability to track the same ROIs over each of the three imaging sessions (Fig. 1E, S1).

First, we examined how population-wide responses to the stimuli changed over days. In the distal apical dendritic ROIs, the difference in responses to unexpected (*A-B-C-U-G*) and expected (*A-B-C-D-G*) sequences increased across days, reaching statistical significance in both L2/3 and L5 by session 3 (Fig. 4A–B, *top*). In contrast, by session 3, the response differences in the somatic ROIs, which were statistically significant in session 1 for L5-S, converged towards zero (Fig. 4A–B, *bottom*). Indeed, specifically comparing the responses to the regular sequence frames (*A-B-C*) and the unexpected frames (*U-G*), we found that the average somatic ROI responses tended to decrease for both expected and unexpected frames over time, though the effect was only statistically significant in L2/3 (Fig. 4A–C, *bottom*). In contrast, in the distal apical dendritic ROIs, we observed an increase in the average responses to the unexpected frames, but not to the regular sequence frames (Fig. 4A–C, *top*). These results indicate that the responses to the unexpected stimuli evolved differently from the responses to the regular sequence frames in these different compartments.

**Figure 4:**
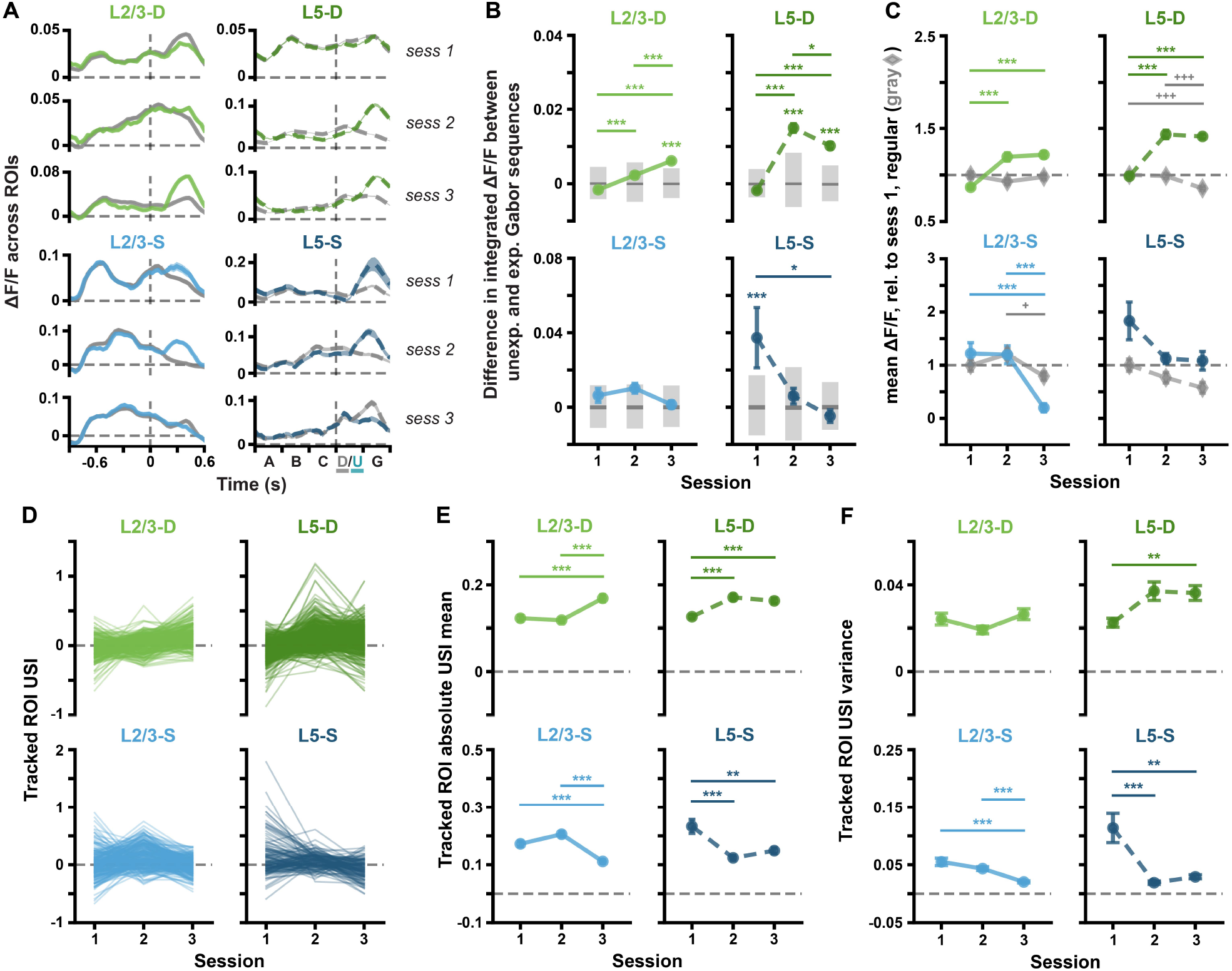
Unexpected Gabor sequences result in different Δ*F/F* and USI changes in different imaging planes. **(A)** Mean (*±* SEM) across ROI mean Δ*F/F* responses to expected (gray, *A-B-C-D-G*) and unexpected (green or blue, *A-B-C-U-G*) Gabor sequences. Dashed vertical lines mark onset of D/U frames. **(B)** Mean (*±* SEM) differences across ROIs in the mean integrated responses to expected vs. unexpected Gabor sequences, as defined in (A). Gray bars show median (dark) and adjusted 95% CIs (light) over randomly expected differences. **(C)** Mean (*±* SEM) across ROIs of the mean Δ*F/F* responses across sequences for regular sequence frames (gray diamonds: *A-B-C*) and unexpected frames (green or blue circles: *U-G*). Responses are calculated relative to session 1 regular responses, marked by dashed horizontal lines. **(D)** Gabor sequence stimulus USIs for all tracked ROIs. Each line represents a single ROI’s USIs over all three sessions. **(E)** Mean (*±* SEM) across the absolute values of the Gabor sequence stimulus USIs for tracked ROIs, as shown in (D). **(F)** Variance (*±* bootstrapped SD) across the Gabor sequence stimulus USIs for tracked ROIs, as shown in (D). *: *p <* 0.05, **: *p <* 0.01, ***: *p <* 0.001 (two-tailed, corrected). ^+^: *p <* 0.05, ^++^: *p <* 0.01, ^+++^: *p <* 0.001 (two-tailed, corrected), for regular stimulus comparisons (gray) in (C). See Table S1 for details of statistical tests and precise p-values for all comparisons.

Importantly, there is evidence that representations in the brain can drift naturally over time, even in the absence of learning [Deitch et al., 2021; Rule et al., 2019]. As such, our above analyses left open the possibility that the changes we observed in the neural responses were not a result of unexpected event-driven learning, but were simply a result of non-specific representational drift. We think this is unlikely because we saw strong directionality to the changes in responses over days that would not be expected from random representational drift: somatic responses to unexpected events decreased towards zero, while distal apical dendritic responses increased across days (Fig. 4A–C).

Nonetheless, to further test for non-specific drift, we also examined the evolution of the responses of the same ROIs to a different, visual flow stimulus (Fig. S5A), which, based on prior work, was unlikely to drive strong expectation violations due to the fact that the visual flow was not coupled to the animals’ movements [Zmarz and Keller, 2016]. In line with this previous work, we observed that although this stimulus drove changes in L2/3-S and L2/3-D, responses in L5-S and L5-D were fairly stable over sessions (Fig. S5B–C) [Jordan and Keller, 2020]. Moreover, in all compartments, the changes in responses to unexpected stimuli and USIs were smaller for the visual flow simuli than the Gabor stimuli (S6A–E). This indicates that our observations of relatively large changes in the responses to the Gabor sequences were stimulus–specific, and hence unlikely to be caused by non-specific representational drift. Altogether, these data support the idea that VisP engages in unsupervised learning in response to unexpected events.

**Figure 5:**
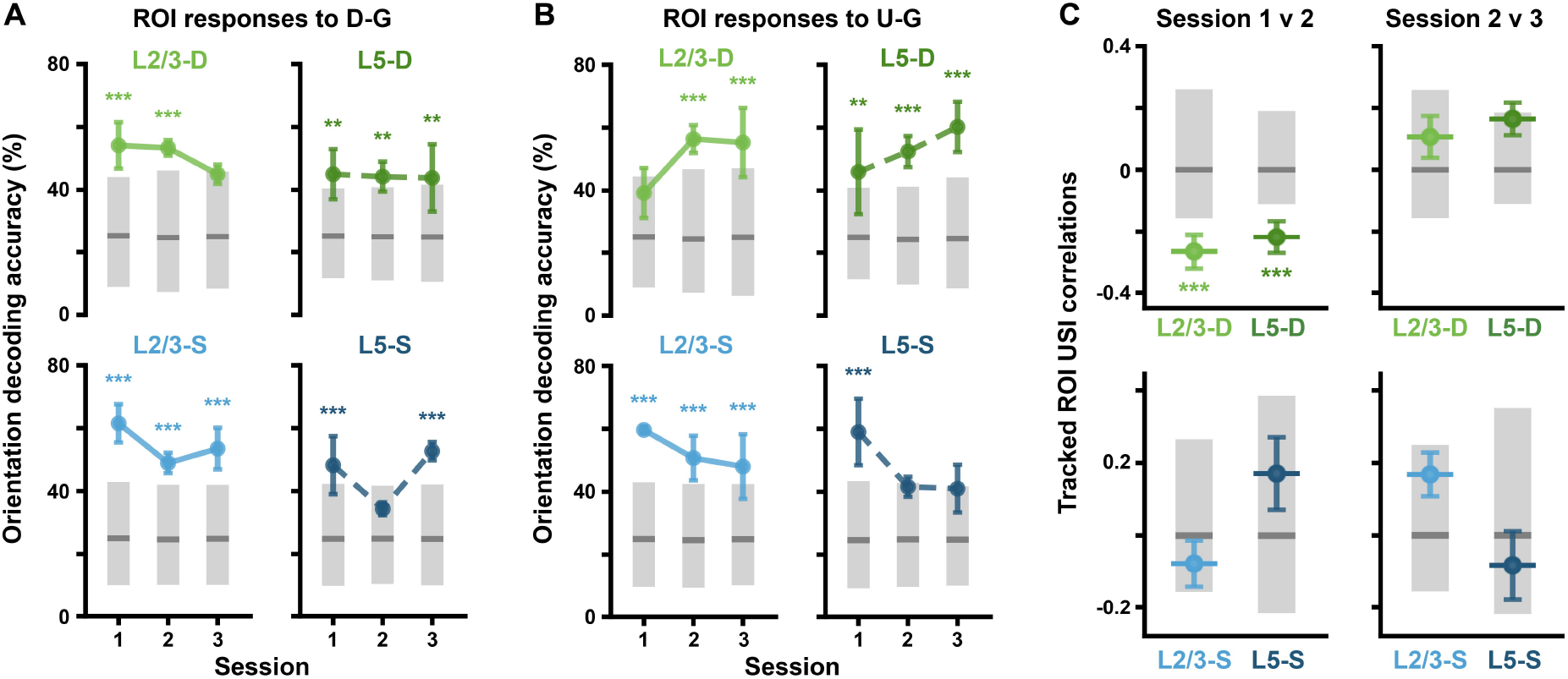
Unexpected Gabor sequences result in predictable Δ*F/F* changes in individual ROIs. **(A)** Balanced accuracy (mean *±* SEM over mice) for classifiers decoding mean Gabor patch orientations from ROI activity during *D-G* frames (2–3 mice per imaging plane, 300 random cross-validation splits per mouse, per session). Gray bars show median (dark) and adjusted 95% CIs (light), computed by shuffling orientation labels. **(B)** Same as (A) but with ROI activity during *U-G* frames. **(C)** USI correlations (*±* bootstrapped SD) between sessions for each plane and session comparison. Gray bars show median (dark) and adjusted 95% CIs (light), computed by shuffling ROI labels. *: *p <* 0.05, **: *p <* 0.01, ***: *p <* 0.001 (two-tailed, corrected, except for (C) where one-tailed (lower), corrected significance is reported). See Table S1 for details of statistical tests and precise p-values for all comparisons.

Given our observations of changes in the responses to the Gabor sequences at the population level, we wondered whether the same effects would be observable for the tracked ROIs. This is important because changes observed in the population-wide responses could, in principle, be driven by only a few ROIs. To test this possibility, we examined the changes over days in the responses of individual ROIs. First, we observed the same patterns as described above when we focused only on the tracked ROIs: i.e., the somatic responses tended to decrease for both regular sequence frames and unexpected frames, whereas the responses to the unexpected frames increased in the distal apical dendrites (Fig. S4B).

Next, in order to understand how the responses to the expected *D* frames versus unexpected *U* frames evolved, we examined the USIs for the tracked ROIs over days. We found that in the somatic compartments, the USIs had converged towards zero over the three sessions (Fig. 4D–F, *bottom*). In contrast, in the distal apical dendritic compartments, the USIs had increased significantly over the three days. This effect was most prominent in L2/3-D (Fig. 4D–E, *top*). These effects were generally consistent across mice (Fig. S4C). Together, these results indicate that individual ROIs altered their responses to the expected and unexpected Gabor sequence stimulus frames over multiple days, in a manner that differed between compartments receiving largely bottom-up inputs (somata) and compartments receiving largely top-down inputs (distal apical dendrites). It is important to note that if these changes were entirely random, then we would not expect to see systematic differences between the somatic and distal apical compartments, as we did here. Instead, this discovery is consistent with the second and third observable signatures of the hierarchical learning hypothesis articulated in the Introduction, since we see changes over days and these differ between the distal apical dendrites and the somata.

### 2.4 Responses to expected and unexpected stimuli change systematically over days

Our preceding analyses showed that neurons in mouse VisP respond differently to expected and unexpected stimuli (Fig. 2), that these responses evolve over days (Fig. 4), and that there is a difference in this evolution between compartments receiving primarily bottom-up or top-down information (Fig. 4). These findings provide evidence for the three observable signatures of hierarchical predictive learning. We next sought to look for evidence that neural circuits use cell-by-cell or distal apical dendritic segment-by-segment differences in responses to expected and unexpected stimuli to guide learning.

A necessary condition for these signals to guide learning is that they contain detailed information about what was unexpected about the stimuli, i.e., information about the orientations of the Gabors. If the neural signals were to contain this information, we should be able to decode the unexpected Gabor patch orientations from the responses to the expected *D* or the unexpected *U* frames. To address this question, we trained linear logistic regression classifiers to identify the mean Gabor patch orientation from the recorded neural responses. Using a cross-validation approach with 300 random splits, we trained the classifiers for each animal and session on 75% of the data, testing them on the remaining (held-out) 25%. We found that in the somatic compartments the classifiers performed significantly above chance for session 1. This performance tended to decrease over sessions until it was at or near chance level by session 3 (Fig. 5A–B, *bottom*). In contrast, in the dendritic compartments, the performance of the decoders started above, or nearly above, chance on session 1 and then improved over sessions for the unexpected *U* frames, but not the expected *D* frames (Fig. 5A–B, *top*). Interestingly, the decoding results for *U* frames paralleled the evolution of the USIs in these compartments. Hence, the signals contain information about the nature of the unexpected orientations in a manner that reflects the extent of differences between expected and unexpected event-driven responses.

We next sought to determine whether the difference in responses to expected and unexpected stimuli was systematic across days at the level of individual ROIs. Specifically, we examined the correlation between ROI USIs in one session and the next. If the USIs do not change systematically over days, the second day’s USI should resemble the first day’s, plus some noise, and hence we should find positive correlations between USIs across days. Conversely, negative correlations between days are evidence of systematic changes, wherein ROIs with the largest USIs on the first day tend to have the smallest USIs on the second day, suggesting a USI-dependent learning mechanism.

To determine whether correlations were significantly different from what would be expected if there was no relationship between an ROI’s USI in one session and the next, we computed null distributions for each imaging plane and session pair by shuffling 10^5^ times the ROI labels within each session. Correlation values below these null distributions were interpreted as reflecting a statistically significant negative correlation between USI values across sessions for individual ROIs.

In the somatic compartments, we found no statistically significant correlations between ROI USIs in one session and the next (Fig. 5C, *bottom*). This suggests that since the overall population tendency is for somatic ROI USIs to converge towards zero over days (Fig. 4D-F, *bottom*), individual ROI USIs on one day are not linearly predictive of their values on a subsequent day. In contrast, ROI USIs in both distal apical dendritic compartments were negatively correlated from session 1 to 2 (Fig. 5C, *top*). As Fig. S7A–B shows, this reflects a tendency for the higher distal apical dendritic ROI USIs to decrease from day 1 to day 2 (*bottom right* quadrants), and for the lower ones to increase even more strongly (*top left* quadrants). ^1^

In contrast, we ran this same analysis on the visual flow stimulus, where changes in representations over sessions were smaller and were only observed in L2/3-D and L2/3-S. Although none of the distal apical dendritic compartments showed USI correlations across days, ROI USIs in somatic compartments were positively correlated across days (Fig. S6F, S7C–D). Given the weak changes in mean ROI USIs across days, these correlations likely reflect a tendency for visual flow USIs to remain constant across days in the somatic compartments.

Altogether, these results demonstrate that at a dendritic segment-by-segment level, the differences between responses to expected and unexpected stimuli change systematically between days. This implicates the distinct responses to expected versus unexpected stimuli in the learning process, as anticipated under a predictive learning hypothesis.

## 3 Discussion

In this study, we explored the question of whether the neocortex learns from unexpected stimuli. This is a central component of a broad class of theories in neuroscience and machine learning that postulate that the brain learns a hierarchical model of the world by comparing predictions about sensory stimuli to the actual stimuli received from the world. This class of theories has several observable signatures in terms of neural responses and how they should evolve over time in response to expected versus unexpected stimuli. We searched for three such observable signatures here, using chronic recordings in mouse VisP, and found evidence in support of each one. First, we observed that neurons responded differently to expected versus unexpected stimuli, which is a precondition for learning from unexpected stimuli. Second, we found that neural responses to the unexpected stimuli changed over days. In contrast, the responses to other stimuli were more stable, suggesting that the unexpected events specifically drove unsupervised learning. Third, the evolution of these responses over days differed between the distal apical dendrites (which are likely driven in large part by top-down feedback from higher-order areas [Budd, 1998; Larkum, 2013a,b]) versus the cell bodies (which are likely driven more by bottom-up sensory input [Budd, 1998; Larkum, 2013a,b]). This indicates that top-down and bottom-up signals are shaped differently by the unsupervised learning process, which is a feature of learning in a hierarchical model. Finally, and going beyond the three main observable signatures, we found that the sensitivity of distal apical dendrites to unexpected events on one day changed systematically by the next day. This final observation shows that changes in activity across days are specific to individual dendritic segments.

Many different forms of hierarchical unsupervised learning have been proposed. The most well-known in neuroscience is probably the predictive coding model of [Rao and Ballard, 1999], along with its variations [Friston and Kiebel, 2009; Spratling, 2017; Whittington and Bogacz, 2017]. But several other models in this vein exist. Examples include Helmholtz machine [Dayan et al., 1995], deep belief net [Hinton and Salakhutdinov, 2006], Bayesian inference [Lee and Mumford, 2003], contrastive learning [Hyvärinen et al., 2019], and contrastive predictive coding [van den Oord et al., 2018] models. What all of these models share is the idea that higher-order association areas make predictions about incoming sensory stimuli, which then get compared to the actual incoming stimuli in order to learn a model of the external world. Hence, all of these models imply the experimental signatures we tested here.

Why do all of these models imply these same observable signatures? First, in order to learn from unexpected events, there must be some available signal that distinguishes unexpected events from expected ones. Thus, a key observable signature is that expected and unexpected events drive distinct responses. Second, all of these models postulate that unexpected events are used to guide unsupervised learning. Thus, stimuli with unexpected components should induce changes in cortical representations. Third, these models all propose that higher-order areas form more abstract representations of the world, and hence the top-down signals communicate something different from the bottom-up signals, which reflect incoming sensory data. Thus, learning should shape these two signals differently, as they encode different aspects of the world. Finally, all of these models propose that the learning algorithm utilizes the difference between expected and unexpected stimuli to shape neural representations. Therefore, our data ultimately provide support for this broad class of models. Future work will attempt to distinguish between the specific models within this broader class.

Based on our data and previous results in the field, we propose a broad conceptual model illustrated in Fig. 6. According to this model, the brain learns an internal representation of the world in associative regions, based on which top-down predictions are provided via the distal apical dendrites to pyramidal neurons in areas like VisP. If incoming stimuli contain unexpected features, i.e., features not predicted at the distal apical dendrites (e.g., unexpected frames in Gabor sequences or an unexpected leaf on an apple, as in Fig. 6, *left*), pyramidal cell somatic and distal apical dendritic activity will reflect the unexpected feature or event. However, with experience, this activity triggers changes to the internal model of the world, such that it better captures the new information provided by the unexpected stimuli (e.g., by accounting for the possibility of different Gabor frames, or of apples with leaves, as in Fig. 6, *right*). As a result, the distal apical dendritic activity becomes more attuned to these novel forms of stimuli.

**Figure 6:**
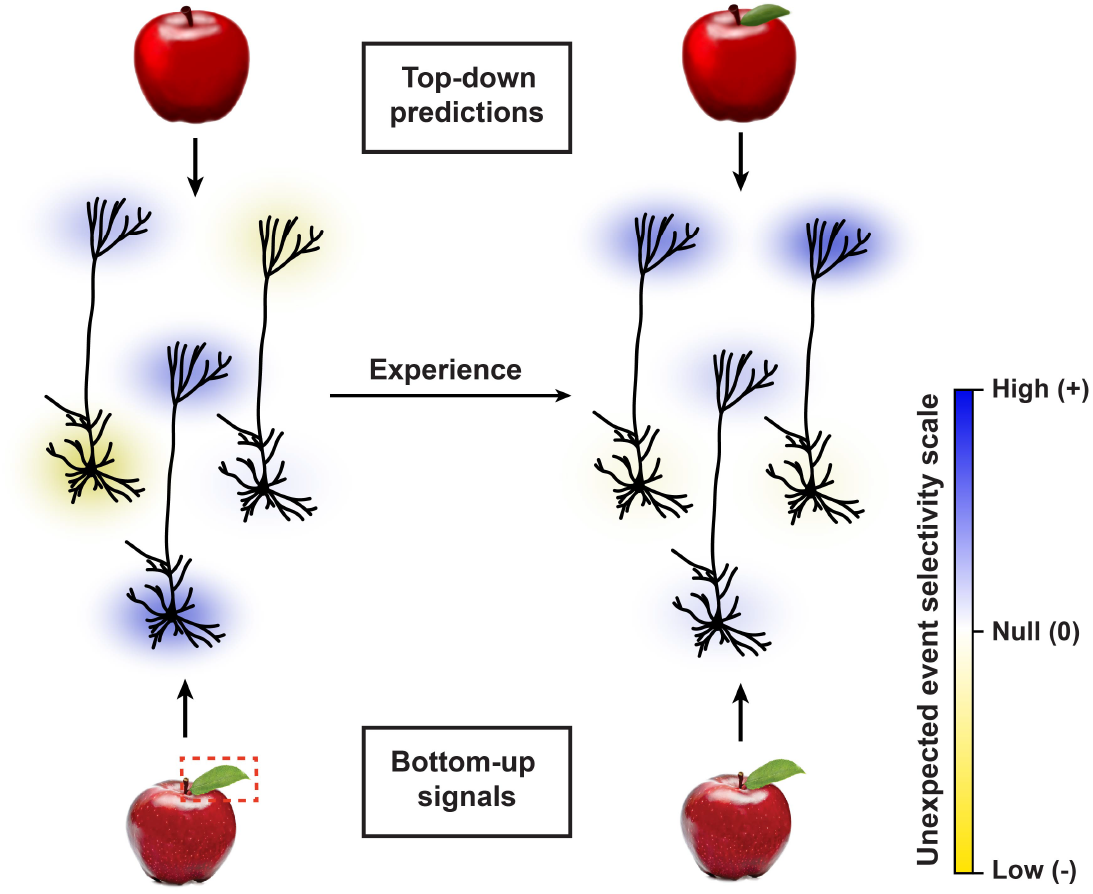
Learning in distal apical dendrites and somata. Illustration of a conceptual model based on our data of how unexpected events drive changes in the neural circuit. With experience, unexpected event selectivity in the somata converges toward 0, whereas it increases overall in the distal apical dendrites, particularly in dendritic segments that initially showed low selectivity.

Notably, our results do not support a simple version of predictive coding wherein excitatory neurons only encode prediction errors. Although the unexpected event responses in the somata did decrease over time, in-line with encoding of errors, the responses in the dendrites increased. This suggests that different computations were reflected in the different compartments of the neurons. Moreover, the finding that the distal apical dendritic signals grow at a population level with exposure to the unexpected stimuli goes counter to proposals implementing predictive coding by using the distal apical dendrite as a site for prediction error calculations [Sacramento et al., 2018; Whittington and Bogacz, 2019]. More experiments with simultaneous imaging of dendrites and cell bodies in different brain regions could help to clarify the distinct computational roles of these neuronal compartments.

There are a number of limitations to this work that must be recognized. First, we were not recording somata and distal apical dendrites in the same neurons. Thus, even though we saw very different evolutions in the responses of the distal apical dendrites and somata to the Gabor sequence stimulus, we cannot say with certainty that these differences hold within individual cells. Indeed, there is data to suggest that coupling between the distal apical dendrites and the somata can sometimes be strong, particularly *in vivo* [Beaulieu-Laroche et al., 2019; Francioni et al., 2019; Larkum et al., 2007]. Nonetheless, other previous work has reported weak coupling between somata and distal apical dendrites [Kerlin et al., 2019; Larkum et al., 2007; Smith et al., 2013], suggesting that this coupling could be context-dependent. Consistent with potential context dependence, we saw clear differences in the evolution of selectivity for unexpected Gabor sequences over time between the somatic and distal apical dendritic compartments, but we did not see these differences in response to the visual flow stimuli (Fig. S6). Since these observations were consistent across mice (Fig. S4C), it seems likely that these results would hold within individual neurons. At the same time, future work using simultaneous multi-plane imaging will be critical to confirm this finding.

Second, though we examined the distal apical dendrites separately from the somata specifically in order to identify potential differences in the processing of top-down and bottom-up inputs, an ideal experiment would record simultaneously from other higher-order brain regions and their projections into visual cortex [Leinweber et al., 2017; Marques et al., 2018]. This would help determine whether the signals we saw in the distal apical dendrites were being calculated locally or in other regions. These experiments would be technically challenging, but they are potentially feasible given recent technical advances in multi-plane mesoscope imaging.

Third, given the nature of our visual stimuli we were unable to measure either the classical receptive fields or the orientation tuning of the neurons. As such, we cannot state with certainty whether these factors could explain the differences in how individual cells responded to expected and unexpected stimuli. However, we observed our results in aggregate across large populations of recorded neurons, presumably with diverse orientation tuning properties and receptive fields. Thus, it is unlikely that idiosyncracies of individual neurons’ orientation selectivities could account for the unexpected event responses. This assertion is supported by our finding that even when we only compared responses for expected and unexpected frames with the same mean orientation we still observed significant differences in the responses. Moreover, we observed significant changes in unexpected event selectivity over days, whereas classic receptive fields and orientation tuning of neurons in mouse VisP are known to be relatively stable over these timescales [Montijn et al., 2016].

Fourth, these experiments were open-loop, and thus did not incorporate any sensorimotor coupling to help shape expectations. On one hand, this is a limitation given that there are a number of reports of apparent sensorimotor predictions and prediction error signals in visual cortex [Keller et al., 2012; Leinweber et al., 2017; Zmarz and Keller, 2016]. On the other hand, the fact that we saw evidence for learning in the open-loop setting suggests that the brain is learning from sensory data alone, in addition to learning from sensorimotor contingencies.

Fifth, and relatedly, our experiments did not incorporate any behavioral training or rewards. It could be the case that the way in which the brain learns from unexpected events is different when those events are relevant to motivated behaviors [Poort et al., 2015]. As such, we cannot say whether the patterns we observed would carry over to task-based learning scenarios.

Finally, it must be recognized that different sensory stimuli, which can present different forms of unexpected events, and recordings in different brain regions may produce different results. To more fully assess the hierarchical predictive learning hypothesis, future work should thoroughly explore the space of possible expected and unexpected sensory stimuli and other regions of the neocortex.

A long-standing goal of neuroscience is to understand how our brains learn from the sensory data that we receive from the world around us. Answers to this question are critical to our understanding of how we build our internal models of the world, and how these govern how we interact with our surroundings. In this work, we monitored changes in the responses of visual cortical neurons in mice while they learned about new external stimuli, and found that these changes were consistent with a broad class of computational models, namely, hierarchical predictive models. Looking forward, we anticipate that these findings could drive substantial progress towards uncovering more specific models describing the brain’s hierarchical predictive learning. To facilitate that progress, our data and analysis software are freely available to other researchers.

## 4 Materials & Methods

### 4.1 Experimental animals and calcium imaging

The dataset used in this paper was collected as part of the Allen Institute for Brain Science’s OpenScope initiative. All animal procedures were approved by the Institutional Animal Care and Use Committee (IACUC) at the Allen Institute for Brain Science. Two transgenic mouse lines (Cux2-CreERT2;Camk2a-tTA;Ai93 and Rbp4-Cre KL100;Camk2a-tTA;Ai93) were used to drive expression of GCaMP6f in layer 2/3 and layer 5 pyramidal neurons, respectively. Mice first underwent cranial window surgery, following which they were housed in cages individually and maintained on a reverse dark-light cycle with experiments conducted during the dark phase. Mice were then habituated over two weeks to head fixation on a running disc, with the visual stimulus presentation being added the second week. Following habituation, they underwent three 70-minute optical imaging sessions within a span of three to six days, with no more than one session occurring per day. Twophoton calcium imaging was performed in the retinotopic center of VisP. Specifically, for each mouse, imaging was performed in either the cell body layer for somatic recordings (175 *µ*m depth for layer 2/3 and 375 *µ*m depth for layer 5) or in cortical layer 1 for distal apical dendritic recordings (50–75 *µ*m depth for layer 2/3 and 20 *µ*m depth for layer 5) across all optical imaging sessions. Sessions that did not meet quality control were excluded from analyses, resulting in 11 mice total (L2/3-D: *n* = 2, L2/3-S: *n* = 3, L5-D: *n* = 3, L5-S: *n* = 3) with three optical imaging sessions each. Full details on the Cre lines, surgery, habituation, and quality control are provided in [de Vries et al., 2020].

Data were collected and processed using the Allen Brain Observatory data collection and processing pipelines [de Vries et al., 2020]. Briefly, imaging was performed with Nikon A1R MP+ two-photon microscopes, and laser excitation was provided at a wavelength of 910 nm by a Ti:Sapphire laser (Chameleon Vision-Coherent). Calcium fluorescence movies were recorded at 30 Hz with resonant scanners over a 400 *µ*m field of view with a resolution of 512 x 512 pixels (Supp. Video 1). Temporal synchronization of calcium imaging, visual stimulation, running disc movement, and infrared pupil recordings was achieved by recording all experimental clocks on a single NI PCI-6612 digital IO board at 100 kHz. Neuronal recordings were then motion corrected, and ROI masks of neuronal somata were segmented as described in [de Vries et al., 2020]. For recordings in layer 1, ROI masks of neuronal dendrites were segmented using the robust estimation algorithm developed by [Inan et al., 2017, 2021], which allows non-somatic shaped ROIs to be identified. This segmentation was run on the motion-corrected recordings, high-pass filtered spatially at 10 Hz and downsampled temporally to 15 Hz. The algorithm parameters were tuned to reject potential ROIs with a peak spatial SNR below 2.5, a temporal SNR below 5, or a spatial corruption index above 1.5, while enabling spatially discontinuous dendritic segments to be identified as part of single ROIs (Fig. S2D). Fluorescence traces for both somatic and dendritic ROIs were then extracted, neuropil-subtracted, demixed, and converted to Δ*F/F* traces, as described in [de Vries et al., 2020; Millman et al., 2020]. Together, neuropil subtraction and the use of a 180-second (5401 sample) sliding window to calculate rolling baseline fluorescence levels *F* for the Δ*F/F* computation ensured that the Δ*F/F* traces obtained were robust to potential differences in background fluorescence between mice and imaging planes. Finally, any remaining ROIs identified as being duplicates or unions, overlapping the motion border or being too noisy (defined as having a mean Δ*F/F* below 0 or a median Δ*F/F* above the midrange Δ*F/F*, i.e., the midpoint between the minimum and maximum) were rejected. In the somatic layers, 15–224 ROIs per mouse per session were identified and retained for analysis, compared to 159–1636 ROIs in the dendritic layers. Lastly, maximum-projection images were obtained for each recording, examples of which are shown in Fig. 1C and E. Briefly, the motion corrected recordings were downsampled to *∼*4 Hz by averaging every 8 consecutive frames, following which the maximum value across downsampled frames was retained for each pixel. The resulting images were then rescaled to span the full 8-bit pixel value range (0–255). Metadata for the dataset is available on GitHub,^2^ and the full dataset is publicly available in Neurodata Without Borders (NWB) format [Ruebel et al., 2019] in the DANDI Archive.^3^

### 4.2 Visual stimulation

During each habituation and imaging session, mice viewed the Gabor sequence stimulus, as well as a visual flow stimulus. The stimuli were presented consecutively for an equal amount of time and in random order. They appeared on a grayscreen background and were projected on a flat 24-inch monitor positioned 10 cm from the right eye. The monitor was rotated and tilted to appear perpendicular to the optic axis of the eye, and the stimuli were warped spatially to mimic a spherical projection screen. Whereas habituation sessions increased in duration over days from 10 to 60 minutes, optical imaging sessions always lasted 70 minutes, comprising 34 minutes of Gabor sequence stimulus and 17 minutes of visual flow stimulus in each direction. Each stimulus period was flanked by one or 30 seconds of grayscreen for the habituation and optical imaging sessions, respectively.

The Gabor sequence stimulus was adapted from the stimulus used in [Homann et al., 2017]. Specifically, it consisted of repeating 1.5-second sequences, each comprising five consecutive frames (*A-B-C-D-G*) presented for 300 ms each. Whereas *G* frames were uniformly gray, frames *A*, *B*, *C*, and *D* were defined by the locations and sizes of the 30 Gabor patches they each comprised. In other words, throughout a session, the locations and sizes of the Gabor patches were the same for all *A* frames, but differed between *A* and *B* frames. Furthermore, these locations and sizes were always resampled between mice, as well as between days, such that no two sessions comprised the same Gabor sequences, even for the same mouse. The location of each Gabor patch was sampled uniformly over the visual field, while its size was sampled uniformly from 10 to 20 visual degrees. Within each repeat of the sequence (*A-B-C-D-G*), the orientations of each of the Gabor patches were sampled randomly from a von Mises distribution with a shared mean and a kappa (dispersion parameter) of 16. This shared mean orientation was randomly selected for each sequence and counterbalanced for all four orientations {0^°^, 45^°^, 90^°^, 135^◦^}. As such, although a large range of Gabor patch orientations were viewed during a session, orientations were very similar within a single sequence. “Unexpected” sequences were created by replacing *D* frames with *U* frames in the sequence (*A-B-C-U-G*). *U* frames differed from *D* frames not only because they were defined by a distinct set of Gabor patch sizes and locations, but also because the orientations of their Gabor patches were sampled from a von Mises distribution with a mean shifted by 90^°^ with respected to the preceding regular frames (*A-B-C*), namely from {90^°^, 135^°^, 180^°^, 225^◦^} (Fig. 2A, Supp. Video 4).

The visual flow stimulus consisted of 105 white squares moving uniformly across the screen at a velocity of 50 visual degrees per second, with each square being 8 by 8 visual degrees in size. The stimulus was split into two consecutive periods ordered randomly, and each defined by the main direction in which the squares were moving (rightward or leftward, i.e., in the nasal-to-temporal direction or vice versa, respectively). Unexpected sequences, or flow violations, were created by reversing the direction of flow of a randomly selected 25% of the squares for 2–4 seconds at a time, following which they resumed their motion in the main direction of flow (Fig. S5A, Supp. Video 5).

Unexpected sequences, accounting for approximately 7% of the Gabor sequences and 5% of visual flow stimulus time, *only* occurred on optical imaging days, and not on habituation days. In particular, each 70-minute imaging session was broken up into approximately 30 blocks, each comprising 30–90 seconds of expected sequences followed by several seconds of unexpected sequences (3–6 seconds for Gabor sequence stimulus and 2–4 seconds for the visual flow stimulus). All durations were sampled randomly and uniformly for each block, across multiples of 1.5 seconds for the Gabor sequence stimulus and of 1 second for the visual flow stimulus.

The stimuli were generated using Python 2.7 [Van Rossum and Drake, 1995] custom scripts based on PsychoPy 1.82.01 [Peirce, 2009] and CamStim 0.2.4, which was developed and shared by the Allen Institute for Brain Science. Code, instructions to reproduce the stimuli, and example videos are available on Github.^4^

### 4.3 Statistical analyses

For most analyses, mean *±* standard error of the mean (SEM) is reported. In cases where the error could not be directly measured over the sample, e.g., the percentage of significant ROI USIs reported in Fig. 2F, a bootstrapped estimate of the error was obtained by resampling the data with replacement 10^4^ times. In these cases, the standard deviation (SD) over the bootstrapped sample is plotted instead, and this is visually signaled by the use of broader error caps (Fig. 2F–G, 4F, 5C, S6B, E–F).

Significance tests, unless otherwise indicated, were computed non-parametrically using permutation tests with 10^5^ shuffles to construct null distributions, based on which confidence intervals (CIs) could be estimated. Where p-values are reported, they are two-tailed (except for Fig. 5C, S6F and S7; see Sec. 4.6 Fluorescence trace analysis, below), and Bonferroni-corrected for multiple comparisons to reduce the risk of Type I errors (false positives). Where 95% CIs are plotted, they are equivalently adjusted using a Bonferroni correction. An exception was made for Fig. 3B, which reports the relationship between the stimuli and behavioral data. Here, Type II errors (false negatives) were considered of greater concern, and thus we reported raw two-tailed p-values in the panel itself. Details of the statistical analyses for all figures, including number of comparisons and corrected p-values, are presented in Table S1.

### 4.4 Running and pupil analysis

Mice were allowed to run freely on a disc while head-fixed during habituation and optical imaging sessions (Fig. 3A, Supp. Video 2). Running information was converted from disc rotations per running frame to cm/s. The resulting velocities were median-filtered with a five-frame kernel size, and any remaining outliers, defined as resulting from a single frame velocity change of at least *±*50 cm/s, were omitted from analyses.

To track pupil diameter during imaging sessions, an infrared LED illuminated the eye ipsilateral to the monitor (right eye), allowing infrared videos to be recorded (Fig. 3A, Supp. Video 3) [Allen Institute for Brain Science, 2017]. We trained a DeepLabCut model from *∼*200 manually labeled examples to automatically label points around the eye, from which we estimated the pupil diameter (*∼*0.01 mm per pixel conversion) [Mathis et al., 2018].^5^ We omitted from analyses outlier frames, defined as resulting from a single-frame diameter change of at least 0.05 mm, which usually resulted from blinking.

Each datapoint in Fig. 3B corresponds to the difference in the mean running velocity or pupil diameter for one block between the unexpected and preceding expected Gabor sequences during session 1, with all blocks being pooled across mice. We computed p-values by comparing the mean difference over all blocks for each plane to a distribution of mean differences, obtained by shuffling the expected and unexpected labels 10^4^ times and calculating the mean difference over all blocks for each shuffle.

### 4.5 ROI tracking across sessions

To track ROIs across days, we employed a custom-modified version of the ROI-matching package developed to track cell bodies across multiple recording days by the Allen Institute for Brain Science [de Vries et al., 2020]. This pipeline implements the enhanced correlation coefficient image registration algorithm to align ROI masks and the graph-theoretic blossom algorithm to optimize the separation and degree of overlap between pairwise matches, as well as the number of matches across all provided sessions [Evangelidis and Psarakis, 2008]. This process produced highly plausible matches for the somatic ROIs; however, it provided some implausible matches for the smaller and more irregularly shaped dendritic ROIs. For the dendritic ROIs, we therefore further constrained the putative matches to those that overlapped by at least 10–20%. Finally, we merged results across all session orderings (e.g., 1-2-3, 1-3-2, 3-1-2), eliminating any conflicting matches, i.e., non-identical matchings that shared ROIs. In total, the modified matching algorithm produced *∼*100–500 highly plausible matched ROIs per plane, i.e., *∼*32–75% of the theoretical maximum number of trackable ROIs (L2/3-D: *n* = 254, L2/3-S: *n* = 261, L5-D: *n* = 516, L5-S: *n* = 129) (Fig. 1E, S1, S8).

### 4.6 Fluorescence trace analysis

For all results except those presented in Fig. 5A–B, S2C, and S4A, C, ROIs were pooled across all mice within an imaging plane for analyses. To enable ROI pooling across mice within imaging planes, each ROI’s Δ*F/F* trace was scaled using robust standardization, i.e., by subtracting the median and then dividing by the interpercentile range spanning the 5*^th^* to 95*^th^* percentile. The only additional exceptions to this are Fig. 4C, S2A-B, S4B, S6A, where unscaled Δ*F/F* traces were used to ascertain how the Δ*F/F* signal itself changed across sessions.

Unexpected event selectivity indices (USIs) were calculated for each ROI separately using Equation 1:

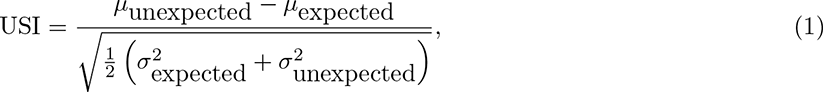

where the means (*µ*_expected_ and *µ*_unexpected_) and variances 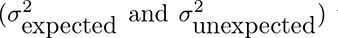 were calculated across integrated Δ*F/F* responses to the expected and unexpected events, respectively. For the Gabor sequences, expected events responses were defined as those spanning *D-G* frames, and unexpected events were defined as those spanning *U-G* frames, with each event therefore spanning 600 ms. Indeed, *G* frames were included in these events, as they did not introduce any new stimuli, but did consistently show persisting ROI responses to *D* or *U* frames (Fig. 2C). For the visual flow stimulus, expected events were defined as the last 2 seconds of expected flow before unexpected flow onset (at which point 25% of the squares reversed direction), while unexpected events were defined as the first 2 seconds of unexpected flow (Fig. S5B). For each ROI, in addition to the true USI, a null distribution over USIs was obtained by randomly reassigning the expected and unexpected event labels to each response 10^4^ times. USIs were deemed significantly low if they lay below the 2.5*^th^* percentile, and significantly high if they lay above the 97.5*^th^* percentile of their null distribution (Fig. 2D).

Note that for Fig. 2G, USIs were calculated using only *D-G* and *U-G* stimuli for which the mean orientations were in {90^°^, 135^◦^}, i.e., the orientations shared by *D* and *U* frames. For each imaging plane, the percentage of significant ROI USIs was then plotted with bootstrapped SDs. Adjusted 95% CIs over chance levels were estimated using the usual approximation method of the binomial CI, with the sample size corresponding to the number of ROIs in the plane (Fig. 2F–G).

For Fig. 4A–B, ROI responses and differences in responses to full expected (*A-B-C-D-G*) and unexpected (*A-B-C-U-G*) sequences were obtained by first taking the mean Δ*F/F* for each ROI across Gabor sequences. Mean Δ*F/F ±* SEM traces were then computed across ROIs and plotted for each session and imaging plane (Fig. 4A). For Fig. 4B, the differences in the traces plotted in (Fig. 4A) were quantified by integrating the mean Δ*F/F* responses over time for each ROI. Mean differences *±* SEM between expected and unexpected sequence responses were then calculated across ROIs and plotted for each session and imaging plane. To further compare ROI responses to regular (*A-B-C*) and unexpected (*U-G*) stimuli, for each ROI, a mean Δ*F/F* was calculated for each set of Gabor frames, and then across sequences (Fig. 4C). The mean Δ*F/F* values thus obtained for each ROI over a given session were then normalized by dividing by the mean Δ*F/F* for regular stimuli across all ROIs from the same mouse in session 1. These normalized means *±* SEM over ROIs were then plotted for each session and plane. Absolute fractional differences between sessions in the responses to unexpected stimuli (Fig. S6B) or in USIs (Fig. S6E) were defined as

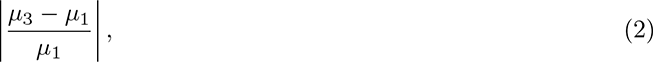

where the subscripts indicate the session over which the mean *µ* is computed. For Fig. S6B, *µ* is the mean of the Δ*F/F* values over all ROIs for the given plane or pooled over all planes, as indicated, for unexpected sequences. As in Fig. 4C, the Δ*F/F* values were calculated relative to the mean expected Δ*F/F* values on session 1 for each mouse. For Fig. S6E, *µ* is the mean of the absolute values of the USIs for the given plane or pooled over all planes for unexpected sequences. Significance tests comparing session results (Fig. 4B–C, E–F, S4B, S5C, S6A, D) and those comparing Gabor sequence and visual flow stimulus results (Fig. S6B, E) were assessed by permuting the session or stimulus labels, respectively, to compute adjusted 95% CIs over results expected by chance.

For the orientation decoding analyses, linear logistic regressions were trained with an *L*_2_ penalty on the multinomial task of classifying the mean Gabor patch orientation for *D-G* frames {0^°^, 45^°^, 90^°^, 135^◦^} or *U-G* frames {90^°^, 135^°^, 180^°^, 225^◦^}. Balanced classifier accuracy was evaluated on the test sets of 300 random cross-validation 75:25 train:test splits of the dataset for each mouse. Importantly, since the *D-G* frame datasets necessarily comprised many more examples than the *U-G* frame datasets (*∼*13x), they were first downsampled for each split to match the number of examples in the corresponding *U-G* frame datasets, thus enabling fairer comparisons between *D-G* and *U-G* classification results. Input data consisted of the Δ*F/F* responses for all ROIs together across *D-G* or *U-G* frames (600 ms). The traces were standardized as described above, but using statistics drawn from the training data only. Mean balanced accuracy across dataset splits was calculated for each mouse, and the mean (*±* SEM) balanced accuracy across mice was plotted for each session and plane. To estimate chance accuracy, shuffled classifier performances were evaluated on 10^5^ random cross-validation dataset splits for each mouse. These classifiers were trained as above, but for each split, the training set orientation targets were shuffled randomly. Null distributions over mean performance were obtained by averaging classifier accuracy for each split across mice, from which adjusted 95% CIs over accuracy levels expected by chance were calculated for each session and plane (Fig. 5A–B).

Pearson correlation coefficients (Fig. 5C, S6F), and the corresponding regression slopes (Fig. S7) were calculated to compare ROI USIs in each imaging plane between sessions. Bootstrapped SDs over these correlations for each plane were then estimated, and adjusted 95% CIs were computed by permuting the ROI labels, such that tracked ROIs were no longer matched together. Here, one-tailed (lower tail) CIs were calculated to identify correlations that were more negative than expected by chance.

## 5 Analysis software

Analyses were performed in Python 3.6 [Van Rossum and Drake, 2009] with custom scripts that are freely available on GitHub,^6^ and were developed using the following packages: NumPy [Harris et al., 2020], SciPy [Jones et al., 2001], Pandas [McKinney et al., 2010], Matplotlib [Hunter, 2007], Scikit-learn 0.21.1 [Pedregosa et al., 2011], and the AllenSDK 1.6.0.^7^ Dendritic segmentation was run in Matlab 2019a [MATLAB, 2019] using the robust estimation algorithm developed by [Inan et al., 2017, 2021]. Pupil tracking was performed using DeepLabCut 2.0.5 [Mathis et al., 2018]. ROIs were matched across sessions using a custom-modified version of the n-way cell matching package developed by the Allen Institute.^8^

## 6 Acknowledgements

The data presented herein were obtained at the Allen Brain Observatory as part of the OpenScope project, which is operated by the Allen Institute for Brain Science. We thank Carol Thompson for her work coordinating the OpenScope project, as well as Christof Koch and John Phillips for their continuous support of the OpenScope project. We thank Wayne Wakeman for data management and support, as well as Nadezhda Dotson, Kiet Ngo and Michael Taormina for their assistance in processing serial two-photon brain sections. We also thank Allan Jones for providing the critical environment that enabled our large-scale team effort. We thank the Allen Institute founder, Paul G. Allen, for his vision, encouragement, and support.

We thank Hakan Inan and Mark Schnitzer, who generously shared with us the code for their robust estimation algorithm [Inan et al., 2017, 2021], and took the time to help us identify the optimal hyperparameter settings for performing dendritic segmentation on the two-photon calcium imaging recordings used in this paper. We also thank Daniel Denman, Stuart Trenholm, and Hubert Banville for helpful feedback on the manuscript.

This work was enabled by the resources provided by Compute Ontario and Compute Canada.^9^

## 7 Author Contributions

These authors contributed equally: CJG, JEP, JAL, as well as BAR, JZ. Experiments were designed by JZ, BAR, TPL, YB. Data was collected by JAL, RA, YNB, SC, PG, IK, EL, JL, KM, CN, TVN, KN, JP, SS, MTV, and AW. Data was analyzed by CJG, JEP, TMH. Supervision was provided by JAL, BAR, JZ, SC, PG, AW. Manuscript was prepared by JEP, CJG, JZ, BAR, TMH.

## 8 Competing Interests

The authors declare no competing interests.

## 9 Funding

This work was supported by the Allen Institute and in part by the Falconwood Foundation. It was also supported by a CIFAR Catalyst grant (JZ and BAR), Canada Research Chair grant (JZ), NSERC Discovery grants (JZ: RGPIN-2019-06379. BAR: RGPIN-2014-04947), Ontario Early Researcher Award (BAR: ER17-13-242), Sloan Fellowship in Neuroscience (JZ), CIFAR Azrieli Global Scholar Award (JZ), Canada CIFAR AI Chair grants (BAR and YB), NSERC Canada Graduate Scholarship - Doctoral Program (CJG), and Ontario Graduate Scholarship (CJG).

## 10 Supplemental Info

### 10.1 Supplemental videos

[Supp. Video 1: Sample two-photon recordings for each imaging plane](https://drive.google.com/file/d/1SkNRvUp67TLc8lRNTgBlq3Yu83Ilw6j1/view?usp=sharing)

This video shows sample two-photon calcium imaging recordings for each of the four imaging planes (L2/3-D, L2/3-S, L5-D, L5-S), which match the maximum-projection images in Fig. 1C. Each recording is from a different mouse and played at 8x the original recording speed.

[Supp. Video 2: Sample of a running recording](https://drive.google.com/file/d/1vR77K1-8cghMBjBLZsahsaIWnQAkaawk/view?usp=sharing)

This video shows a sample recording of a mouse running on a disc, during stimulus presentation, on an optical imaging day.

[Supp. Video 3: Sample of an annotated pupil recording](https://drive.google.com/file/d/1veGb8F_zG_661BJgYkxq4Dj3L2J_oUn2/view?usp=sharing)

This video shows a sample recording of a mouse pupil, during stimulus presentation, on an optical imaging day. The right pupil, ipsilateral to the stimulus presentation screen, is shown. It is annotated with tracking markers which are inferred by the DeepLabCut model and used to measure pupil diameter. Specifically, the small, filled blue dots mark the 8 tracked pupil poles, and the yellow ellipse marks the elliptical pupil shape inferred from the tracked poles.

[Supp. Video 4: Gabor sequence stimulus example](https://drive.google.com/file/d/1qqirJp9fFdOhbYhyfjRpfosiS11_kBEe/view?usp=sharing)

This video shows example expected and unexpected sequences for the Gabor sequence stimulus. As described in the Materials & Methods, each frame lasts 300 ms, resulting in 1.5-second sequences. Within each expected sequence (*A-B-C-D-G*), all Gabor patches share a mean orientation, sampled from {0^°^, 45^°^, 90^°^, 135^◦^}. Within each unexpected sequence (*A-B-C-U-G*), *U* frame Gabor patches are shifted by 90^°^ with respect to the rest of the sequence’s mean Gabor patch orientation. In this example video, each frame is labelled at the bottom right. Additionally, expected sequences are signalled with a green circle, and unexpected sequences, with a red circle. None of these annotations appeared during the actual experiments, when the animals viewed the stimulus.

[Supp. Video 5: Visual flow stimulus example](https://drive.google.com/file/d/1YJKYYmvgmdMwuSe2qv2Tk4jg3PVtmqTp/view?usp=sharing)

This video shows example expected and unexpected sequences for the visual flow stimulus. Example sequences are shown for temporal to nasal (leftward), followed by nasal to temporal (rightward) main flow. As described in the Materials & Methods, during unexpected flow, 25% of the squares temporarily reverse their direction of flow.

### 10.2 Supplemental figures

**Figure S1:**
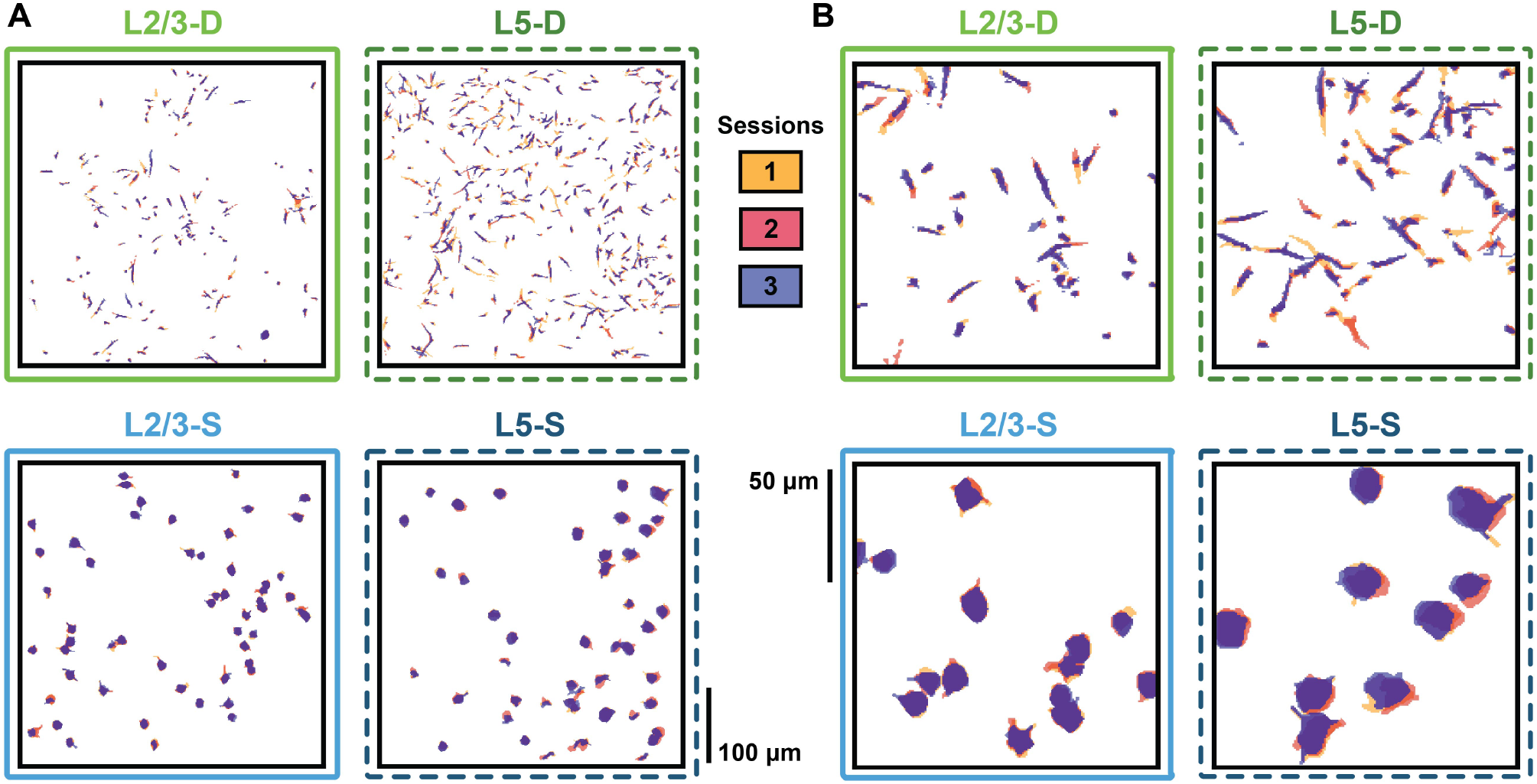
ROIs are successfully tracked in each plane. **(A)** Full field of view mask overlays for ROIs tracked across all three sessions for an example mouse in each plane. **(B)** Enlarged views from (A) showing individual tracked ROI overlays for each plane. The tracking pipeline reliably produced highly plausible ROI matches across all three sessions in each imaging plane.

**Figure S2:**
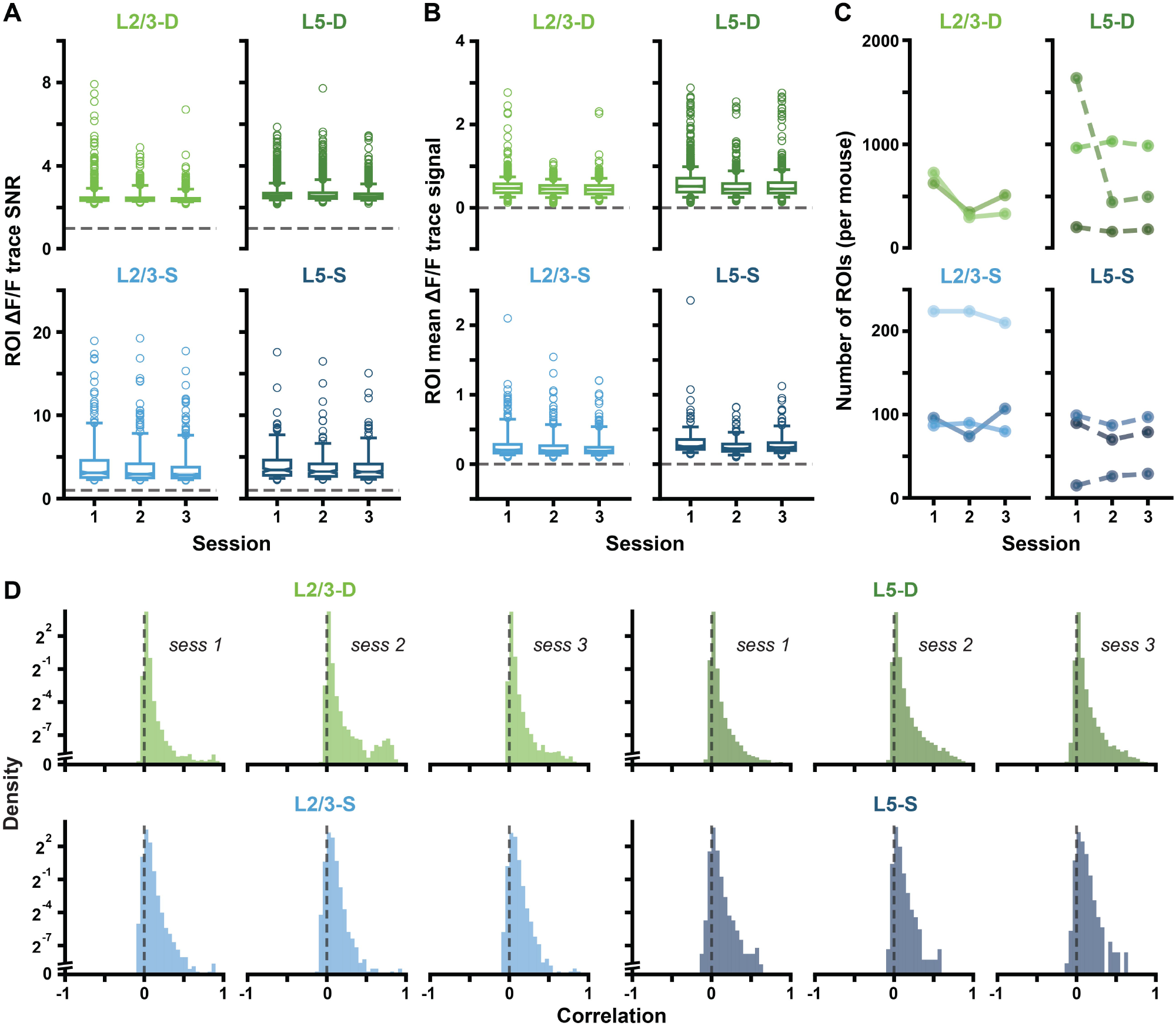
ROI SNR, signal and numbers are consistent across sessions. **(A)** Δ*F/F* trace SNRs for each ROI. For each session and plane, boxplots show the medians of the ROI SNR distributions, as well as the 25*^th^* to 75*^th^* percentiles, with the whiskers extending from the 5*^th^* to 95*^th^* percentiles. SNR was calculated for each ROI as follows. First, parameters (mean, SD) of a normal distribution over noisy activity were estimated based on the lower tail of the ROI’s full activity distribution. The 95*^th^* percentile of the parameterized noise distribution was then defined as that ROI’s noise threshold. ROI SNRs were then calculated as the ratio between their mean activity above the noise threshold (signal), and the SD of their parameterized noise distribution. Dashed horizontal lines mark 1, i.e., noise level. SNR levels were consistent across sessions within imaging planes. **(B)** Mean Δ*F/F* trace signal, where each datapoint corresponds to an ROI. Boxplots drawn as in (A), and signal is defined as described in (A). Signal levels were consistent across sessions within imaging planes. **(C)** The number of ROIs was generally stable across sessions for each mouse, except one in L5-D. **(D)** Distributions of pairwise ROI correlations, plotted on a log scale. The log scale is linearized near 0, as signalled by the axis break. Pairwise correlations were computed over full session fluorescence traces, which were smoothed using a four-point moving average. In all sessions, lines and planes, the vast majority of the correlation mass was concentrated near 0. The log scale reveals that the small amount of mass remaining is distributed similarly between lines and planes, largely below 0.5.

**Figure S3:**
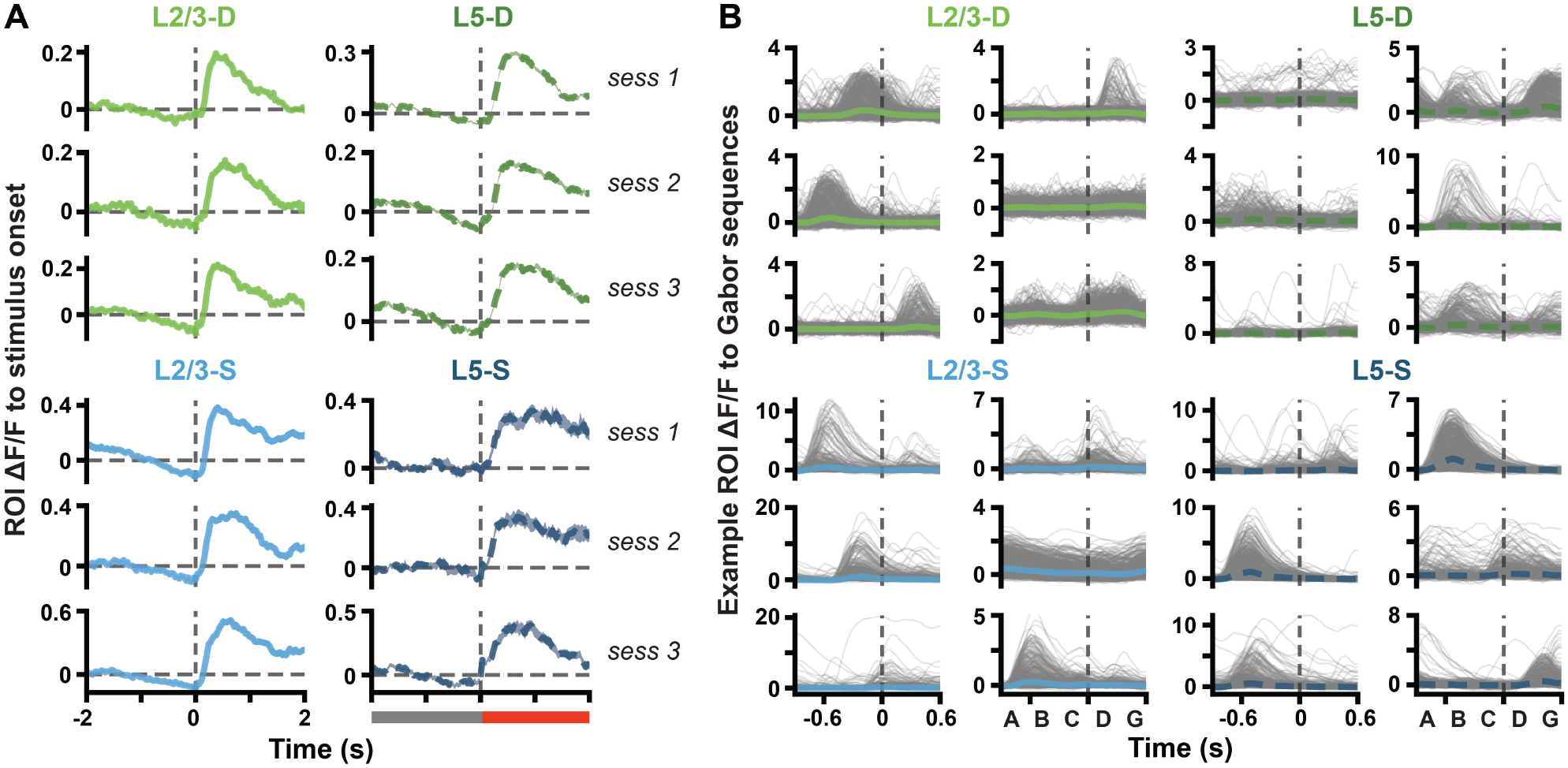
ROIs are responsive to stimulus onset, and to Gabor sequences. **(A)** Mean (*±* SEM) Δ*F/F* response traces across ROI mean responses to stimulus onset (Gabor sequence or visual flow) from grayscreen. Dashed vertical line at time 0 marks stimulus onset, also signalled by the gray bar becoming red (bottom of right column). In all planes and sessions, ROI populations show clear responses to stimulus onset. **(B)** Δ*F/F* response traces to each expected Gabor sequence (gray) for example ROIs. Mean (*±* SEM) Δ*F/F* responses across sequences are plotted in blue or green. Dashed vertical lines mark onset of D frames. Plotted ROIs were randomly selected from session 1 ROIs deemed consistently responsive to Gabor sequences, based on the following criteria: (1) their SNR was above the median for the session, (2) the median pairwise correlation between their individual sequence responses, as well as the SD and skew of their mean response, were each above the 75*^th^*percentile for the session. As in Fig. S2D, responses to individual sequences were smoothed using a four-point moving average, for correlation calculation and plotting, only. In each imaging plane, numerous ROIs were found which were responsive to various components of the Gabor sequences.

**Figure S4:**
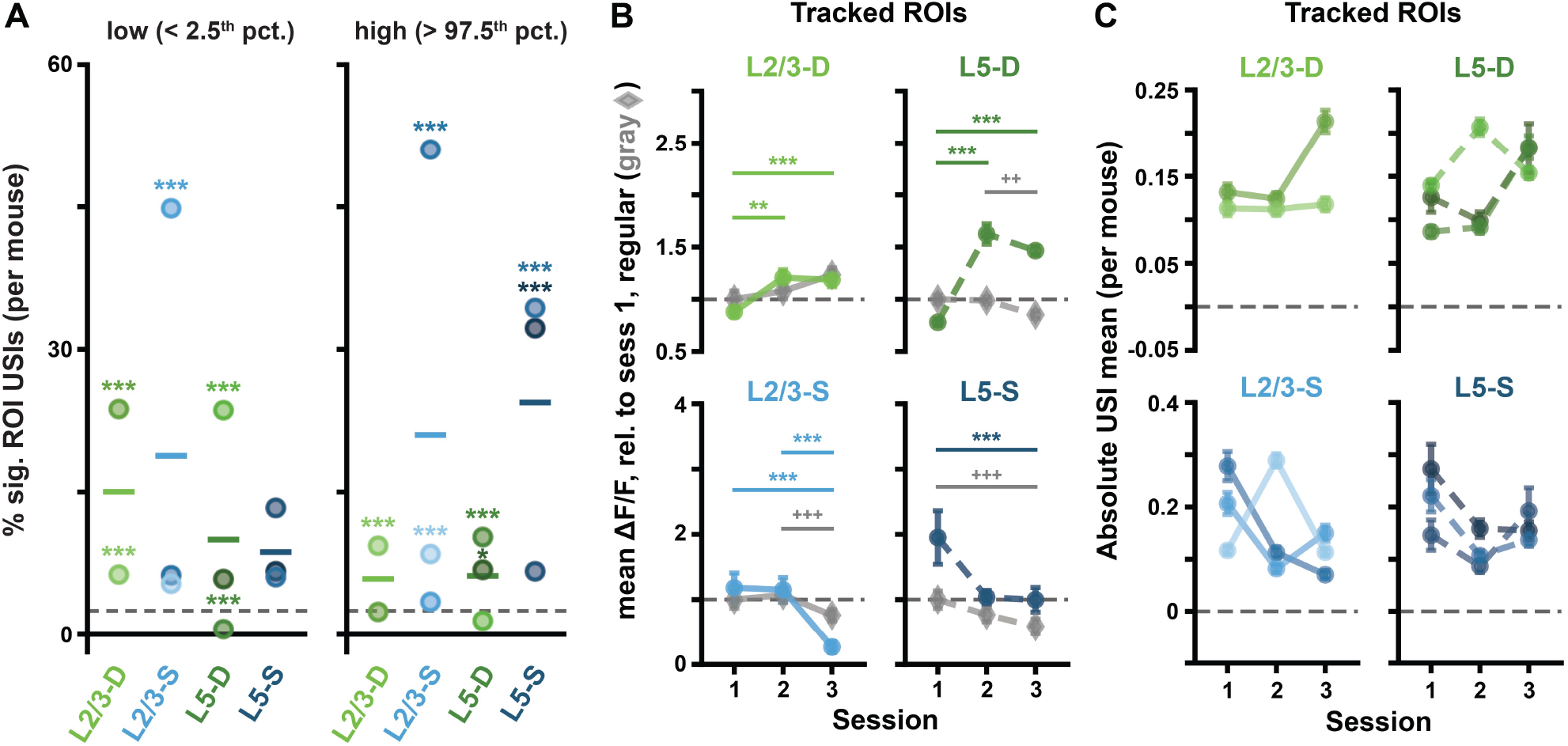
ROI responses to unexpected Gabor sequences are consistent in tracked ROIs and across mice. **(A)** Percentage of significant USIs in session 1 for each plane, where each dot corresponds to a separate mouse. Significance for each datapoint was evaluated against its own adjusted binomial CI (not shown). Lines show the pooled percentage for each plane, as plotted in Fig. 2F. Dashed horizontal lines mark the theoretical chance level (2.5%). Results are consistent with those pooled across mice, with 10 out of the 11 animals showing a higher percentage of significant ROI USIs than expected by chance in at least one tail (2F). **(B)** Mean (*±* SEM) across tracked ROIs of the mean Δ*F/F* responses across sequences for regular sequence frames (gray diamonds: *A-B-C*) and unexpected frames (green or blue circles, *U-G*), as in Fig. 4C. Responses are calculated relative to session 1 regular responses, marked by dashed horizontal lines. Results are consistent with the full ROI population results (Fig. 4C). **(C)** Mean (*±* SEM) across the absolute values of the Gabor sequence stimulus USIs for tracked ROIs, as in Fig. 4E, but split by mouse. Results are consistent with those pooled across mice (Fig. 4E). *: *p <* 0.05, **: *p <* 0.01, ***: *p <* 0.001 (two-tailed, corrected). ^+^: *p <* 0.05, ^++^: *p <* 0.01, ^+++^: *p <* 0.001 (two-tailed, corrected), for regular stimulus comparisons (gray) in (B). See Table S1 for details of statistical tests and precise p-values for all comparisons.

**Figure S5:**
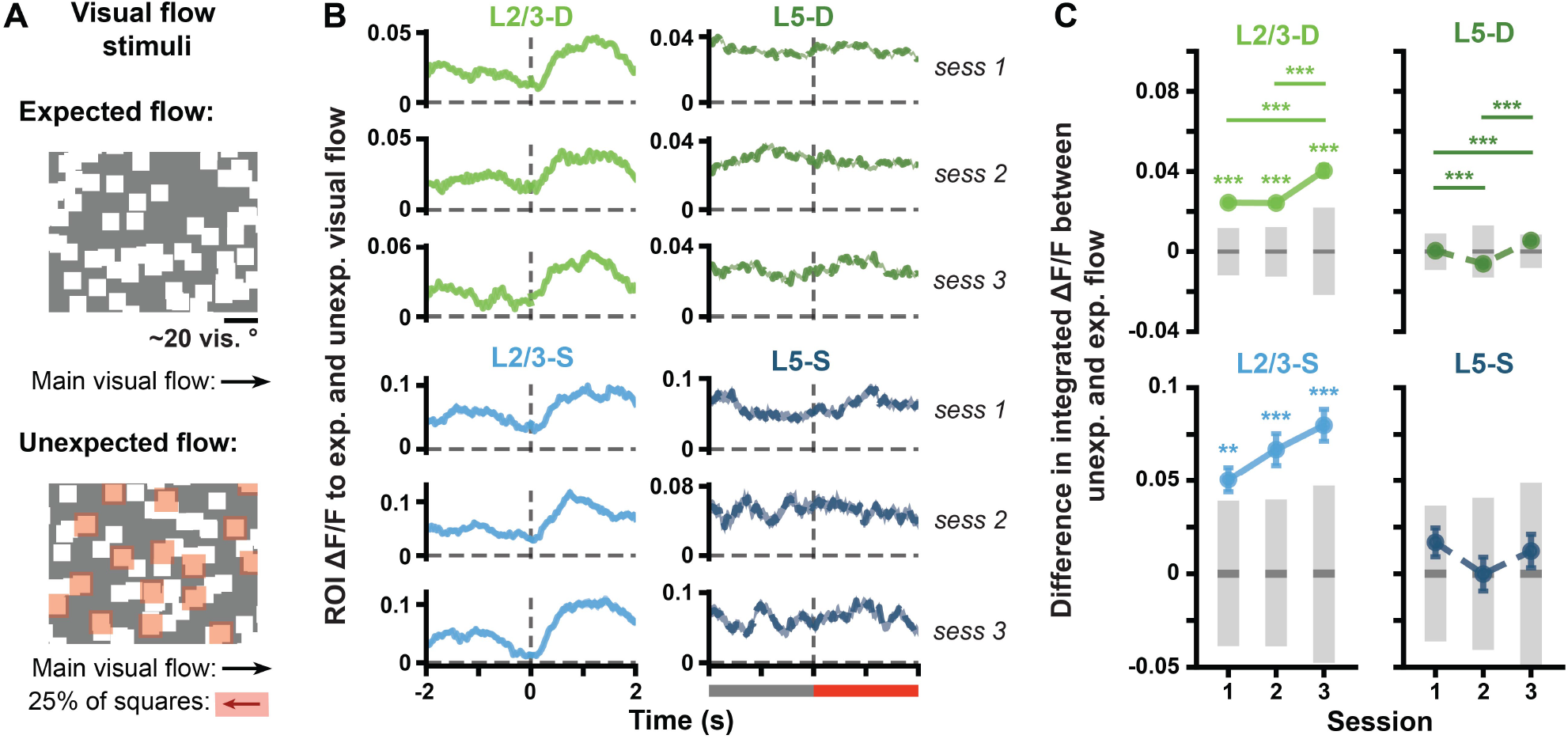
L2/3 ROIs respond to the onset of unexpected visual flow. **(A)** Visual flow stimulus. Squares moved together at the same velocity across the screen during expected flow (*top*). At random times (unexpected flow, *bottom*), 25% of the squares, highlighted here in red for illustrative purposes, reversed direction for 2–4 seconds (see Materials & Methods, and Supp. Video 5). **(B)** Mean (*±* SEM) across ROI mean Δ*F/F* responses to visual flow sequences. Expected and unexpected visual flow sequences were defined as for the USI calculation, namely over the 2 seconds preceding unexpected visual flow onset and following its onset, respectively. Dashed vertical line at time 0 marks the onset of unexpected visual flow, also signalled by the gray bar becoming red (bottom of right column). **(C)** Mean (*±* SEM) differences across ROIs in the mean integrated responses to expected vs. unexpected visual flow, as defined in (B). Gray bars show median (dark) and adjusted 95% CIs (light) over randomly expected differences. Whereas the L5 somatic and distal apical dendritic populations did not respond significantly differently to expected vs. unexpected flow, both L2/3 somatic and distal apical dendritic populations showed a significant difference in responses, which increased over days in the dendrites. These findings are consistent with recent work by [Jordan and Keller, 2020] showing that L2/3 neurons integrate visual flow mismatch information, whereas L5 neurons do not appear to do so. *: *p <* 0.05, **: *p <* 0.01, ***: *p <* 0.001 (two-tailed, corrected). See Table S1 for details of statistical tests and precise p-values for all comparisons.

**Figure S6:**
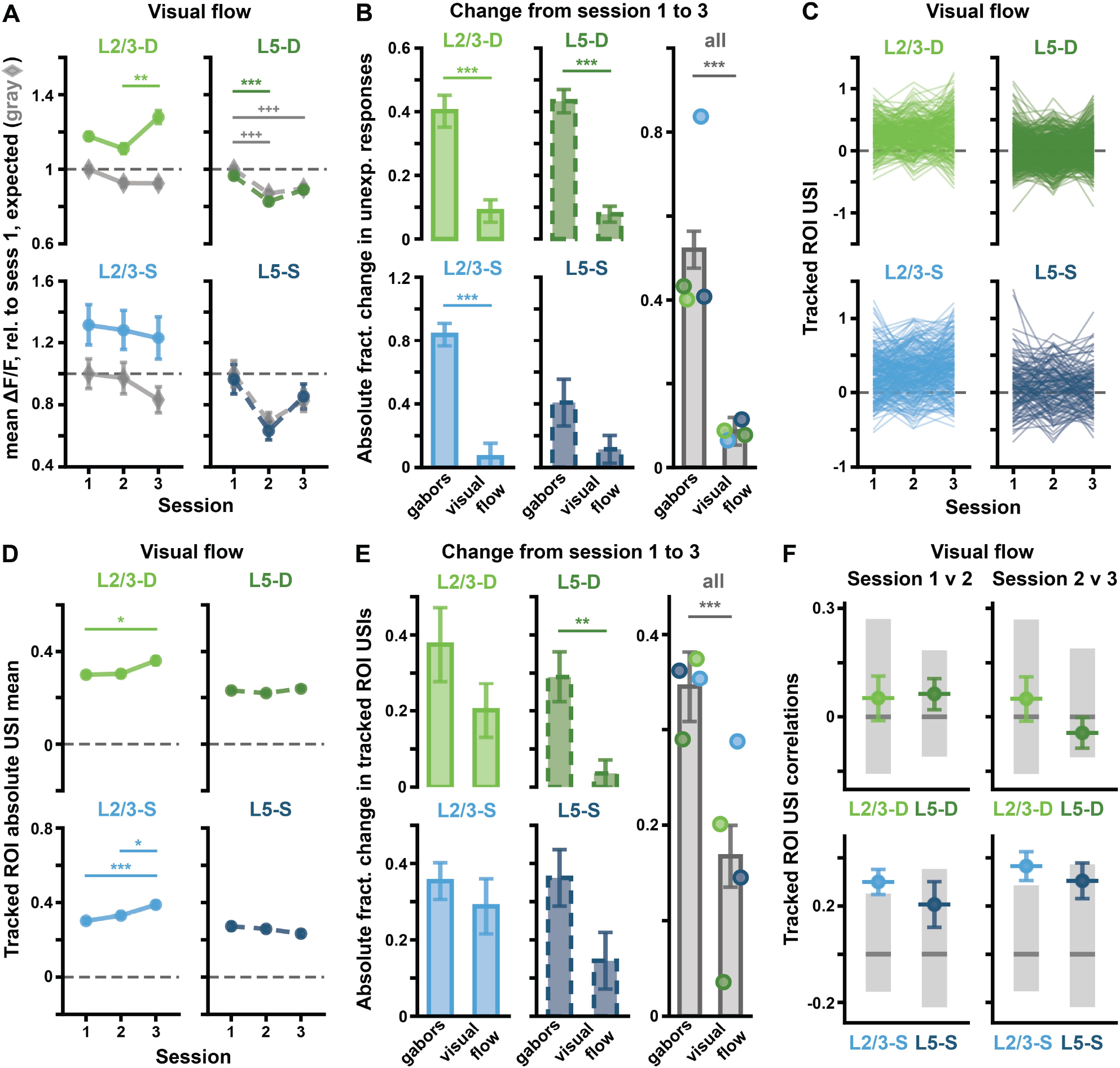
Unexpected visual flow sequences do not result in the same Δ*F/F* changes across sessions as unexpected Gabor sequences do. **(A)** Mean (*±* SEM) across ROIs of the mean Δ*F/F* responses across sequences for expected flow (gray diamonds) and unexpected flow (green or blue circles), as defined in Fig. S5B. Responses are calculated relative to session 1 expected responses, marked by dashed horizontal lines. Corresponds to Fig. 4C for Gabor sequences. **(B)** Absolute fractional change (*±* bootstrapped SD) in mean unexpected responses from session 1 to 3 for Gabor sequence vs. visual flow stimulus for each plane (*left* and *middle* columns), and pooled across all planes (*right* column) (see Equation 2). In all imaging planes except L5-S, changes in ROI responses to unexpected stimulus from session 1 to 3 were significantly greater for the Gabor stimulus than for the visual flow stimulus. **(C)** Visual flow stimulus USIs for all tracked ROIs. Each line represents a single ROI’s USIs over all three sessions. Corresponds to Fig. 4D for Gabor sequences. **(D)** Mean (*±* SEM) across the absolute values of the visual flow stimulus USIs for tracked ROIs, as shown in (C). Corresponds to Fig. 4E for Gabor sequences. **(E)** Similar to (B), but here mean (*±* bootstrapped SD) absolute fractional changes in USIs from session 1 to 3 across tracked ROIs are plotted (see Equation 2). In L5-D and all compartments combined, changes in USIs for tracked ROIs from session 1 to 3 were significantly greater for the Gabor stimulus than for the visual flow stimulus. **(F)** Correlations (*±* bootstrapped SD) for each plane and session comparison. Gray bars show median (dark) and adjusted 95% CIs (light), computed by shuffling ROI labels. Corresponds to Fig. 5C for Gabor sequences. Unlike the Gabor sequence stimulus, only positive correlations are observed for the visual flow stimulus, and they are in the somatic compartments instead of the dendritic ones. *: *p <* 0.05, **: *p <* 0.01, ***: *p <* 0.001 (two-tailed, corrected, except for (F) where one-tailed (lower), corrected significance is reported). ^+^: *p <* 0.05, ^++^: *p <* 0.01, ^+++^: *p <* 0.001 (two-tailed, corrected), for expected stimulus comparisons (gray) in (A). See Table S1 for details of statistical tests and precise p-values for all comparisons.

**Figure S7:**
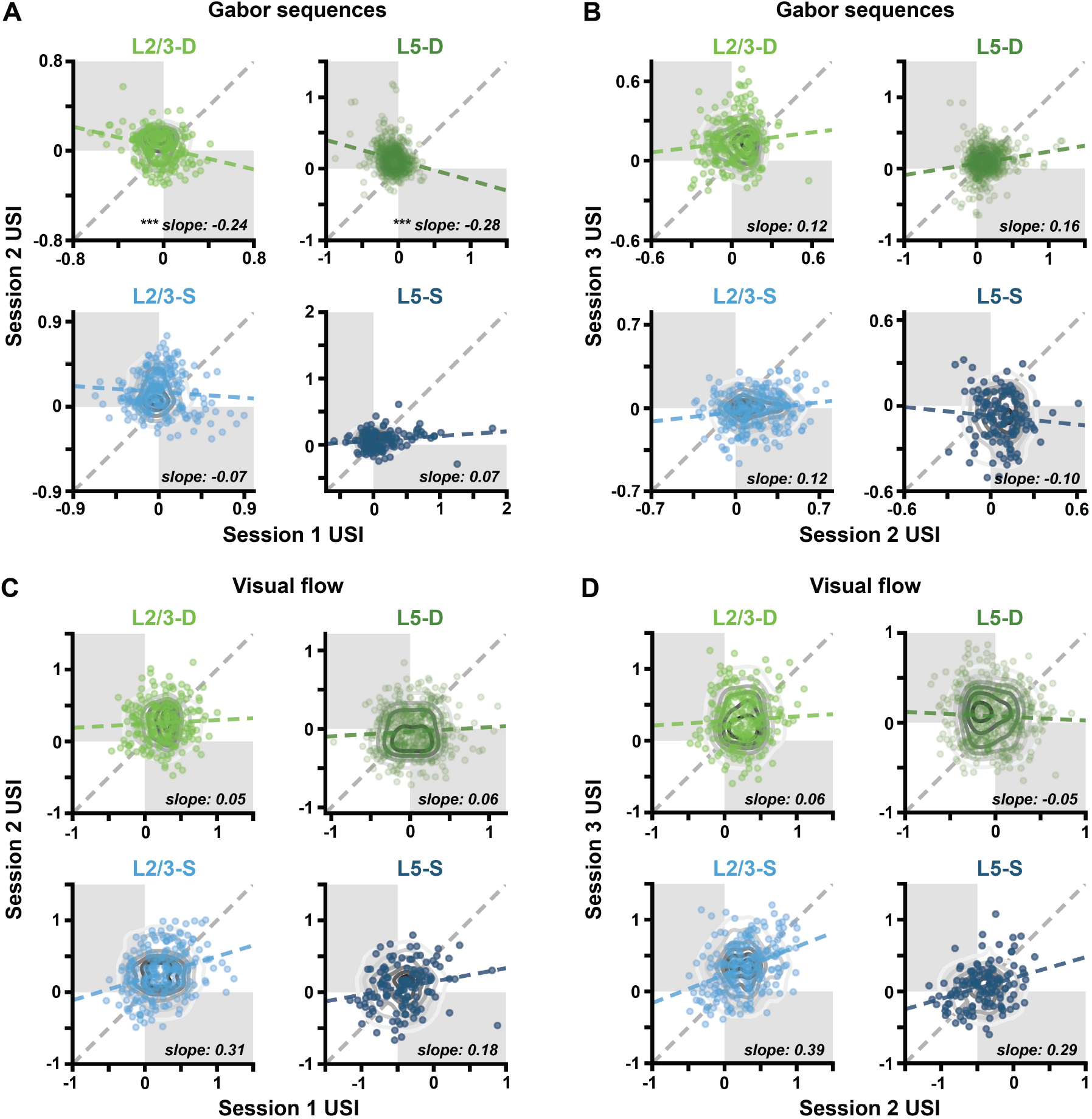
Gabor sequence USIs are negatively correlated between sessions 1 and 2 in the distal apical dendrites. **(A)** Gabor USI scatterplots showing correlations between sessions 1 and 2. Each point reflects a single tracked ROI’s USIs on two sessions. Gray contour lines show null distributions, computed by shuffling ROI labels, as in Fig. 5C. The estimated regression slopes for each plane (blue or green, dashed) are plotted against the identity line (gray, dashed). Opposite quadrants are shaded in gray. Significance markers next to reported slope values correspond to the correlation significance testing results reported in Fig. 5C, and S6F. **(B)** Same as in (A), but for Gabor sequence USIs in sessions 2 and 3. **(C)** Same as in (A), but for visual flow USIs in sessions 1 and 2. **(D)** Same as in (A), but for visual flow USIs in sessions 2 and 3. Only L2/3-D and L5-D ROI Gabor sequence USIs show significant negative correlations. These correlations are between the USIs of sessions 1 and 2, and have slopes of -0.24 and -0.28, respectively. *: *p <* 0.05, **: *p <* 0.01, ***: *p <* 0.001 (one-tailed (lower), corrected). See Table S1 for details of statistical tests and precise p-values for all comparisons. Specifically, for (A–B), see the entries for Fig. 5C, and for (C–D), see the entries for Fig. S6F.

**Figure S8:**
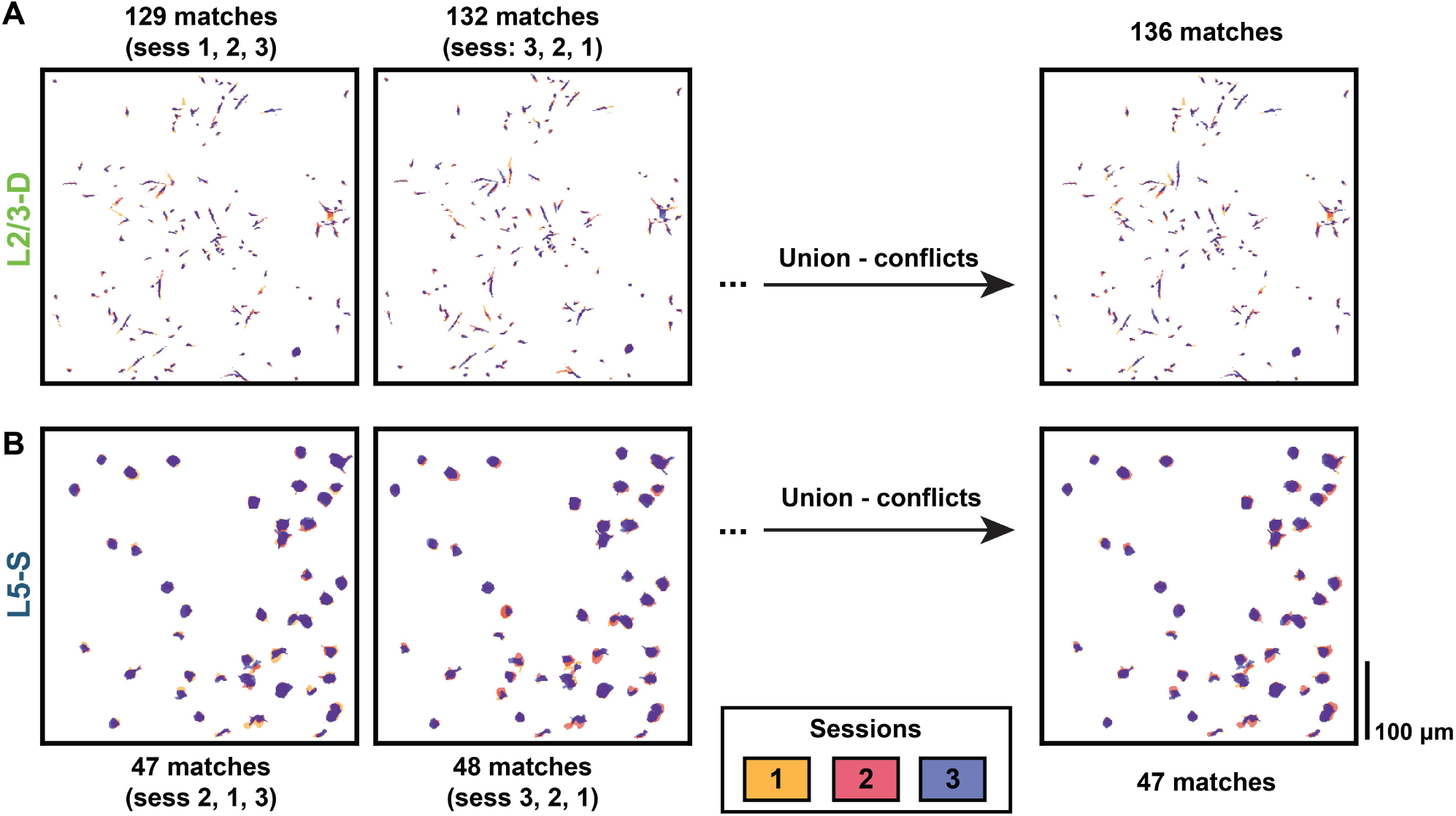
Dendritic ROI matches vary more with session ordering than somatic ROI matches do. **(A)** Example L2/3-D mouse with ROIs matched across sessions. The order in which the session images are aligned slightly affects which ROIs are matched. (*Left*) Permutation with the smallest number of matched ROIs. (*Middle*) Permutation with the largest number of matched ROIs. (*Right*) Taking the union of matches across all session permutations while removing conflicting matches (matches comprising at least one ROI that also appears in a different match) enables the quantity and quality of matched ROIs to be optimized. In this example, four pairwise matches were identified as conflicts and removed, yielding 136 final matches. **(B)** Same as (A), but for a L5-S mouse. The variation in number of matched ROIs across session orderings for somata was generally far less than that for dendrites due to their larger sizes and more regular shapes. Combining matched ROIs across all permutations did nonetheless, in this example mouse, enable two of the pairwise matches to be identified as conflicts and removed, yielding 47 final matches.

### 10.3 Supplemental table

**Table S1:**
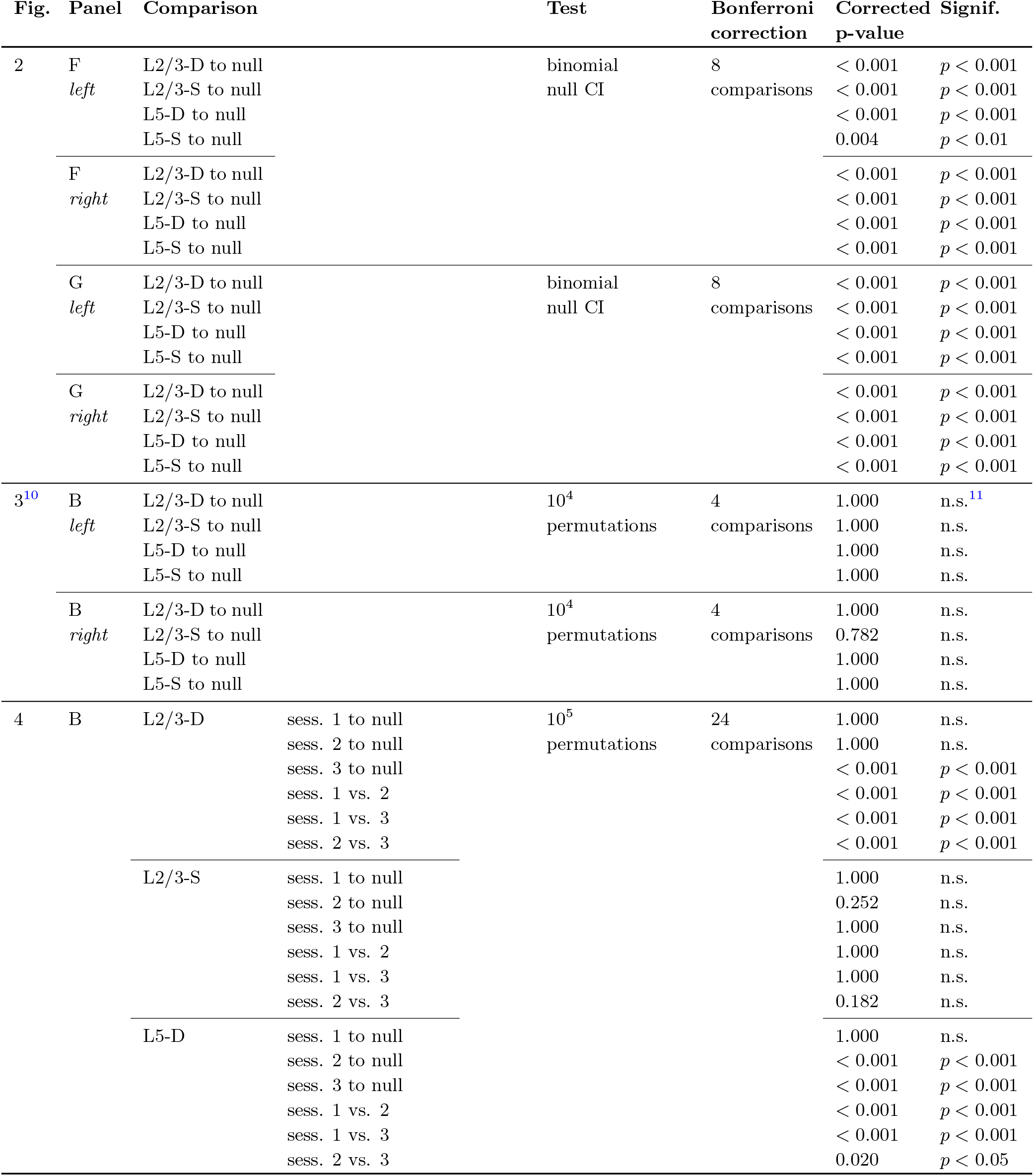
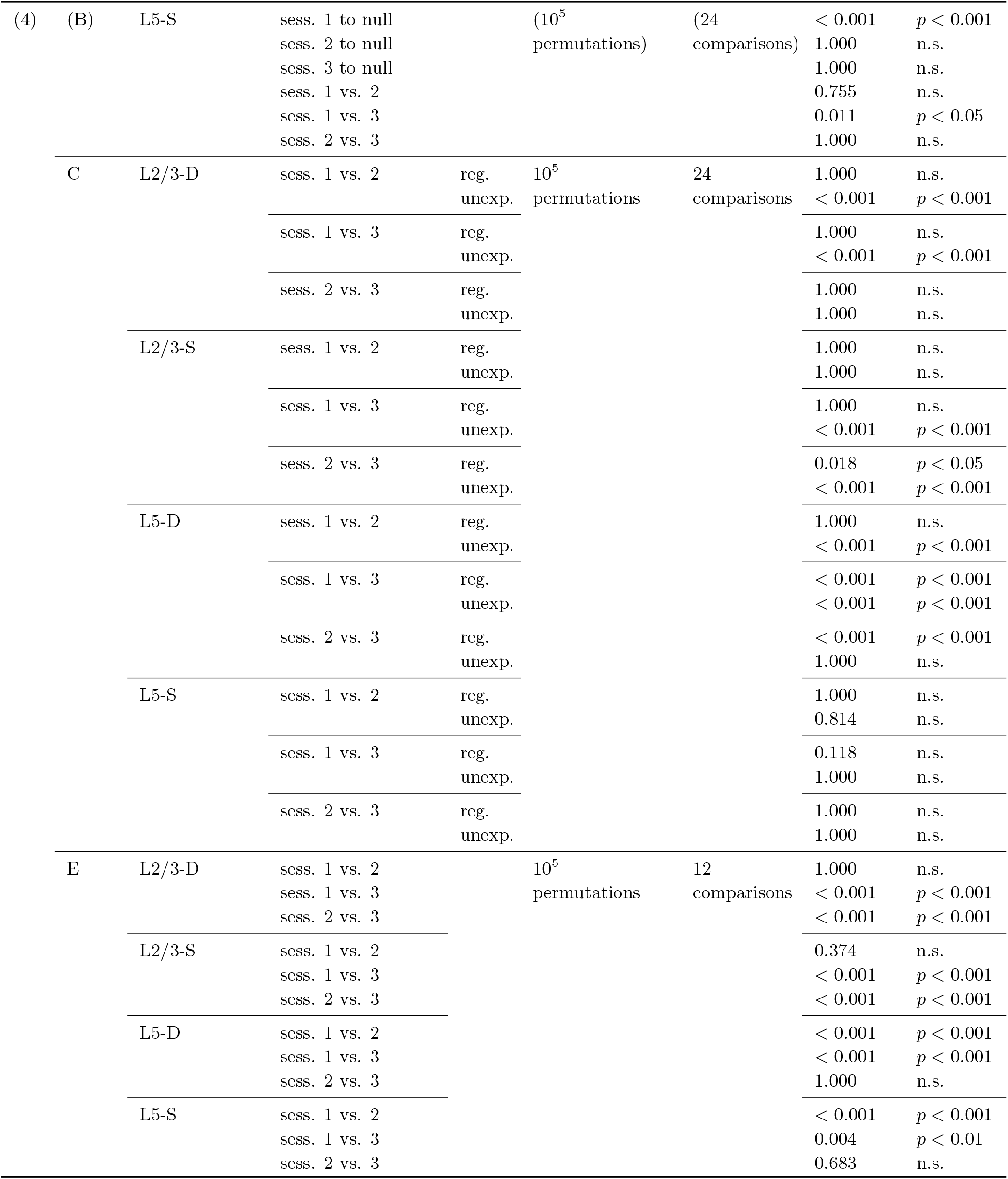
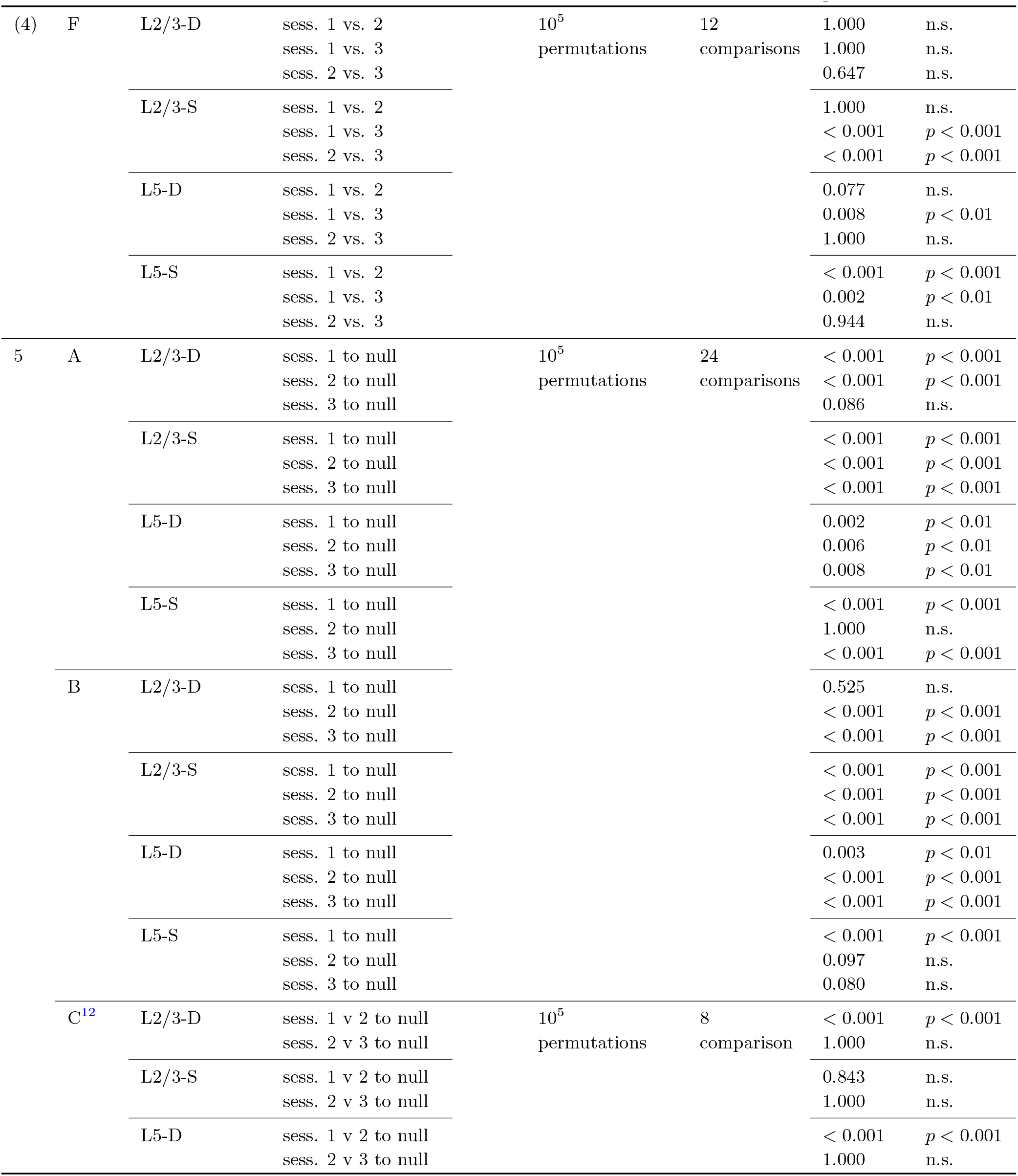
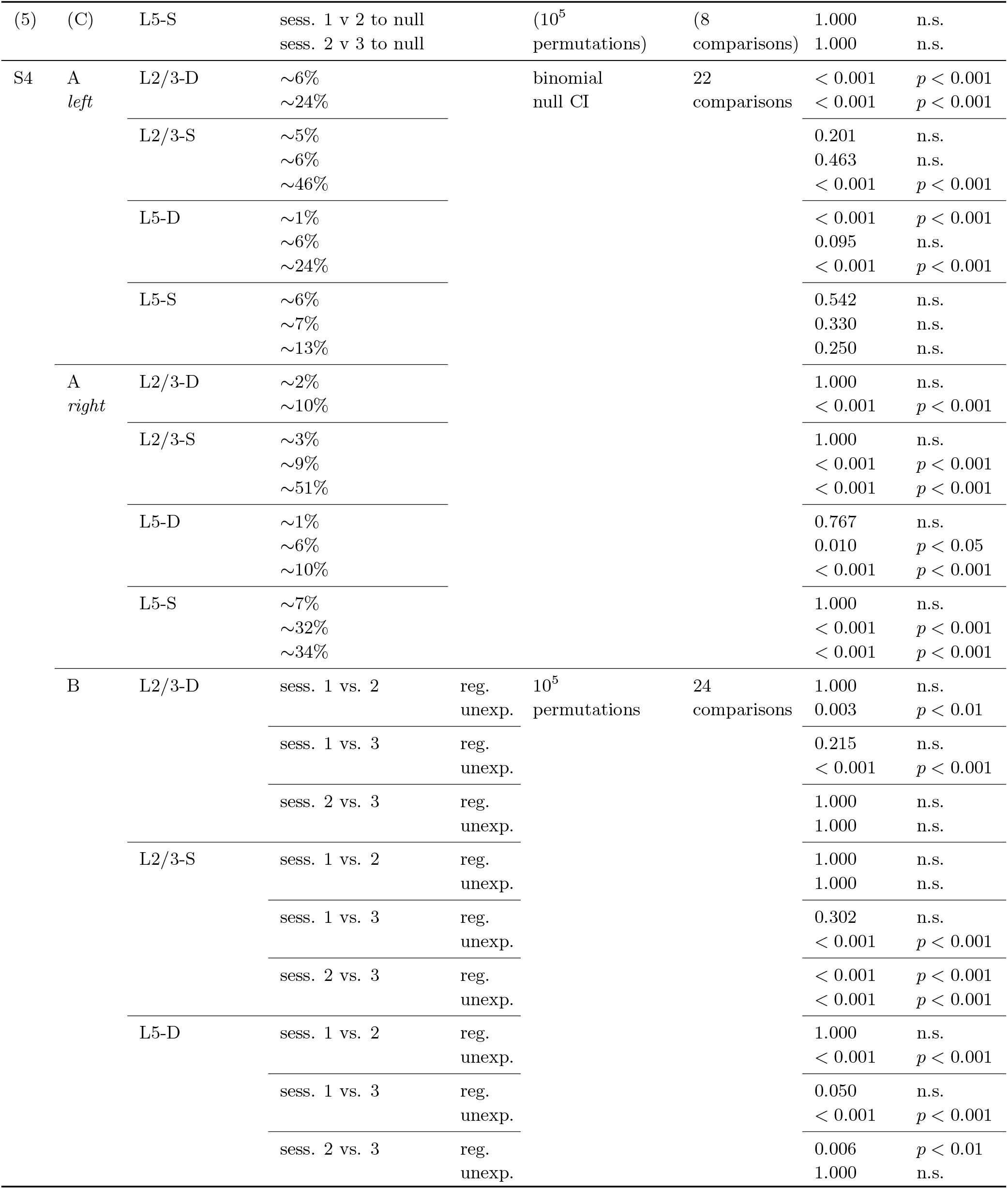
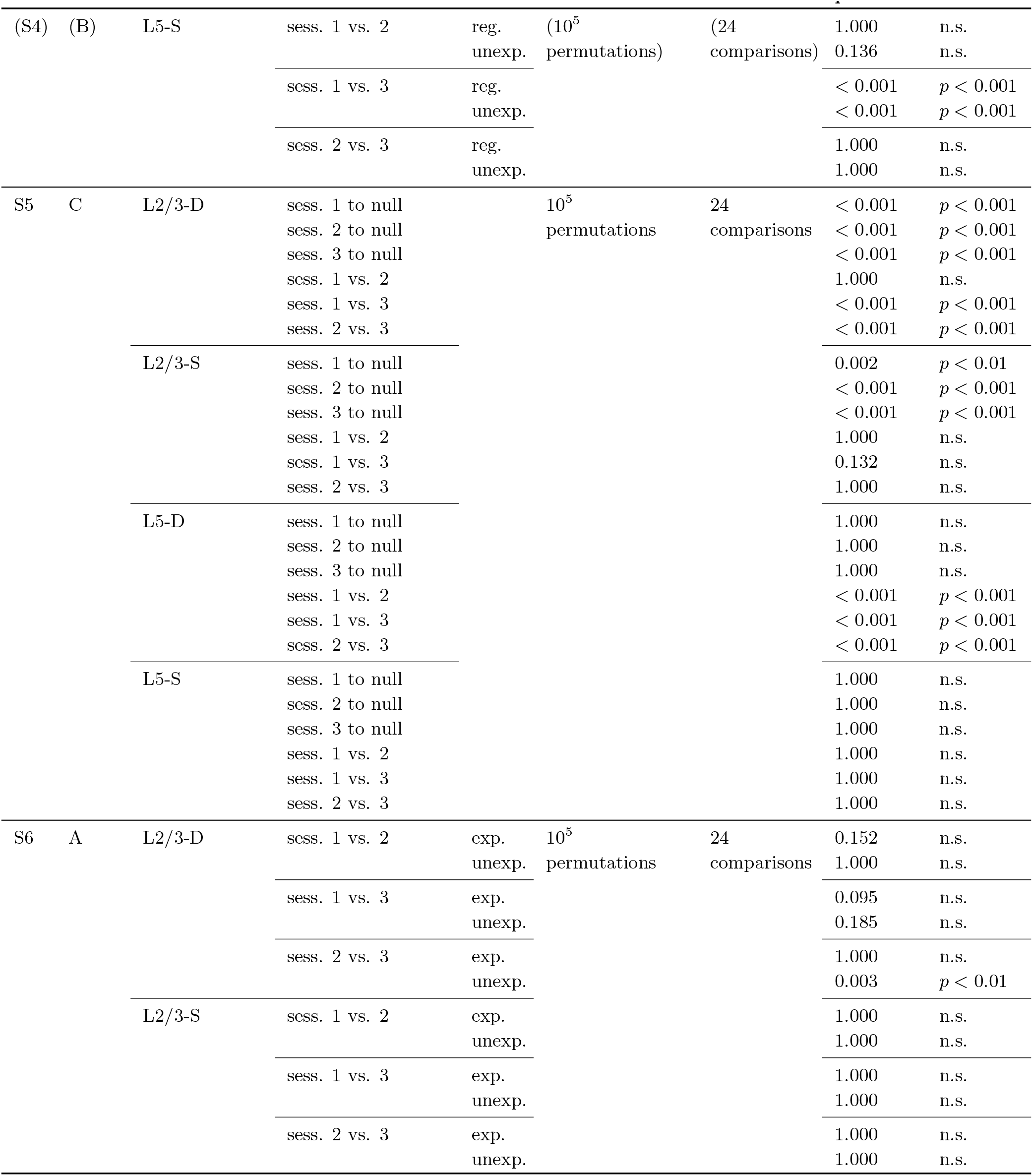
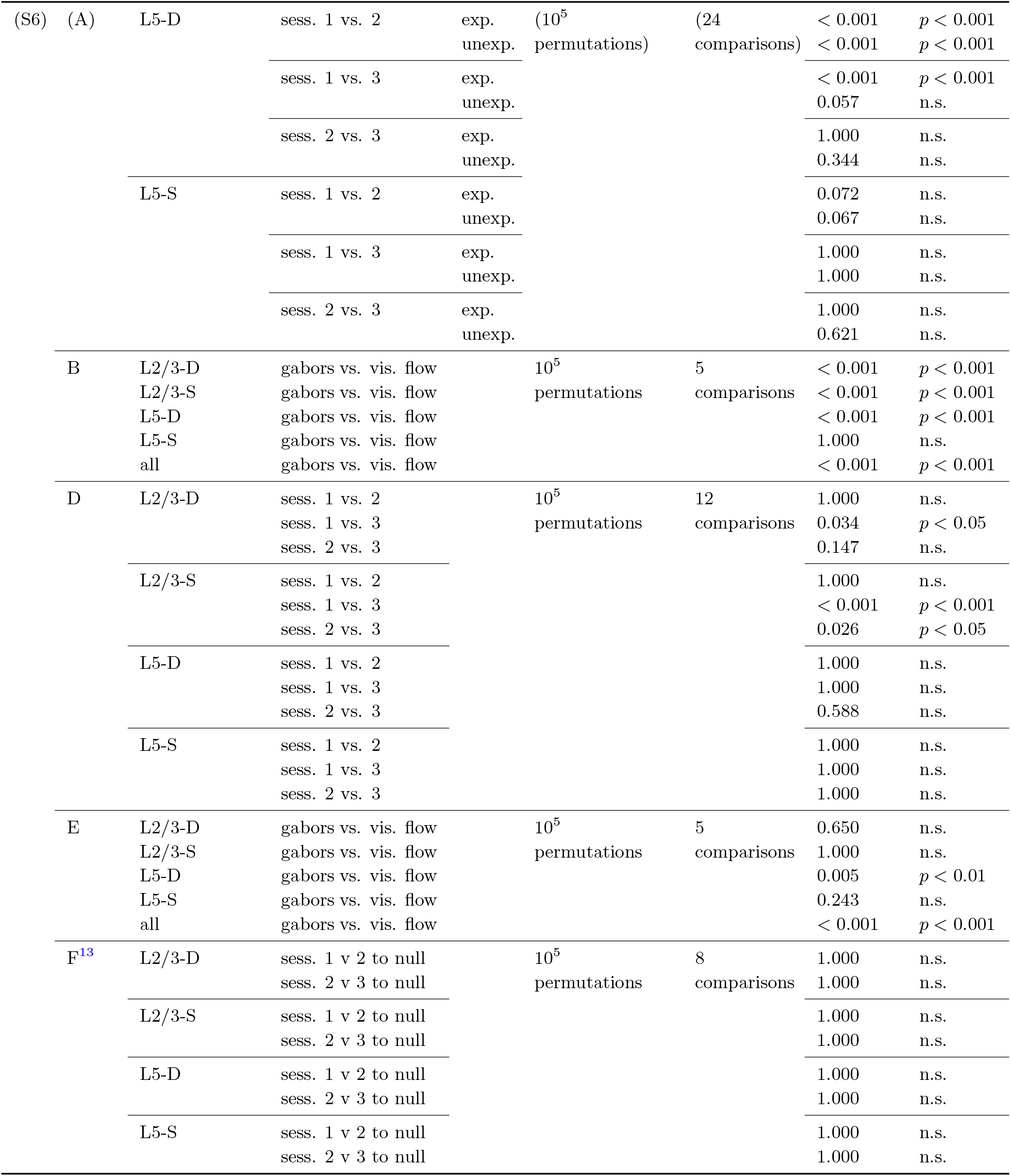
Summary of statistical tests and results for all figures.

## Footnotes

1 The previous version of this paper on *bioRxiv* showed different correlation results, which we have revised here in three ways. 1) Generally, we added more imaging sessions to our analyses to improve robustness. 2) For Fig. 5C and S6F, in order to specifically assess learning at the individual ROI level, we replaced session permutations with ROI permutations. 3) In light of these new results, we updated our interpretations of changes at the level of individual ROIs.

2 https://github.com/jeromelecoq/allen_openscope_metadata/tree/master/projects/credit_assignement

3 https://gui.dandiarchive.org/#/dandiset/000037

4 https://github.com/colleenjg/cred_assign_stimuli

5 https://allensdk.readthedocs.io/en/latest/allensdk.internal.brain_observatory.eye_calibration.html

6 https://github.com/colleenjg/OpenScope_CA_Analysis

7 https://github.com/AllenInstitute/AllenSDK

8 https://github.com/AllenInstitute/ophys_nway_matching/tree/main/nway

9 www.computeontario.ca and www.computecanada.ca

10 In contrast to the p-values reported directly in Fig. 3, the p-values reported here are corrected for multiple comparisons.

11 n.s.: not significant

12 One-tailed t-tests were used here (lower tail).

13 One-tailed t-tests were used here (lower tail).

## Notes

https://gui.dandiarchive.org/\#/dandiset/000037

https://github.com/colleenjg/OpenScope_CA_Analysis

### Competing Interest Statement

The authors have declared no competing interest.

### Summary of Updates

The license has been changed to CC-BY.

## References

Allen Institute for Brain Science (2017). Visual coding overview. Technical report, http://observatory.brain-map.org/visualcoding/.

Bakhtiari, S., Mineault, P. J., Lillicrap, T., Pack, C. C., and Richards, B. A. (2021). The functional specialization of visual cortex emerges from training parallel pathways with self-supervised predictive learning. In Advances in Neural Information Processing Systems, volume 34, pages 2901–2910.

Beaulieu, C. and Cynader, M. (1990). Effect of the richness of the environment on neurons in cat visual cortex. I. Receptive field properties. Developmental Brain Research, 53(1):71–81.

Beaulieu-Laroche, L., Toloza, E., Brown, N. J., and Harnett, M. T. (2019). Widespread and highly correlated somato-dendritic activity in cortical layer 5. Neuron, 103(2):235–241.

Budd, J. M. (1998). Extrastriate feedback to primary visual cortex in primates: a quantitative analysis of connectivity. Proceedings of the Royal Society of London. Series B: Biological Sciences, 265(1400):1037–1044.

Chen, T., Kornblith, S., Norouzi, M., and Hinton, G. (2020). A simple framework for contrastive learning of visual representations. In International Conference on Machine Learning, volume 37, pages 1597–1607.

Christensen, E. and Zylberberg, J. (2020). Models of the primate ventral stream that categorize and visualize images. bioRxiv:2020.02.21.958488.

Dayan, P., Hinton, G. E., Neal, R. M., and Zemel, R. S. (1995). The Helmholtz machine. Neural Computation, 7(5):889–904.

de Vries, S. E., Lecoq, J. A., Buice, M. A., Groblewski, P. A., Ocker, G. K., Oliver, M., Feng, D., Cain, N., Ledochowitsch, P., Millman, D., et al. (2020). A large-scale standardized physiological survey reveals functional organization of the mouse visual cortex. Nature Neuroscience, 23(1):138–151.

Deitch, D., Rubin, A., and Ziv, Y. (2021). Representational drift in the mouse visual cortex. Current Biology, 31(19):4327–4339.

Devlin, J., Chang, M.-W., Lee, K., and Toutanova, K. (2018). BERT: Pre-training of deep bidirectional transformers for language understanding. arXiv preprint arXiv:1810.04805.

Evangelidis, G. D. and Psarakis, E. Z. (2008). Parametric image alignment using enhanced correlation coefficient maximization. IEEE Transactions on Pattern Analysis and Machine Intelligence, 30(10):1858–1865.

Fiser, A., Mahringer, D., Oyibo, H. K., Petersen, A. V., Leinweber, M., and Keller, G. B. (2016). Experience-dependent spatial expectations in mouse visual cortex. Nature Neuroscience, 19(12):1658–1664.

Francioni, V., Padamsey, Z., and Rochefort, N. L. (2019). High and asymmetric somato-dendritic coupling of V1 layer 5 neurons independent of visual stimulation and locomotion. Elife, 8.

Friston, K. and Kiebel, S. (2009). Predictive coding under the free-energy principle. Philosophical Transactions of the Royal Society B: Biological Sciences, 364(1521):1211–1221.

Garrido, M. I., Kilner, J. M., Stephan, K. E., and Friston, K. J. (2009). The mismatch negativity: a review of underlying mechanisms. Clinical Neurophysiology, 120(3):453–463.

Grill, J.-B., Strub, F., Altché, F., Tallec, C., Richemond, P., Buchatskaya, E., Doersch, C., Avila Pires, B., Guo, Z., Gheshlaghi Azar, M., et al. (2020). Bootstrap your own latent - a new approach to self-supervised learning. In Advances in Neural Information Processing Systems, volume 33.

Harris, C. R., Millman, K. J., van der Walt, S. J., Gommers, R., Virtanen, P., Cournapeau, D., Wieser, E., Taylor, J., Berg, S., Smith, N. J., et al. (2020). Array programming with NumPy. Nature, 585(7825):357–362.

Hawkins, J. and Blakeslee, S. (2004). On Intelligence. Macmillan.

Higgins, I., Stringer, S., and Schnupp, J. (2017). Unsupervised learning of temporal features for word categorization in a spiking neural network model of the auditory brain. PLoS One, 12(8):e0180174.

Hinton, G. E. and Salakhutdinov, R. R. (2006). Reducing the dimensionality of data with neural networks. Science, 313(5786):504–507.

Homann, J., Koay, S. A., Glidden, A. M., Tank, D. W., and Berry, M. J. (2017). Predictive coding of novel versus familiar stimuli in the primary visual cortex. bioRxiv:197608.

Huang, L., Ledochowitsch, P., Knoblich, U., Lecoq, J., Murphy, G. J., Reid, R. C., de Vries, S. E., Koch, C., Zeng, H., Buice, M. A., et al. (2021). Relationship between simultaneously recorded spiking activity and fluorescence signal in gcamp6 transgenic mice. Elife, 10:e51675.

Hunter, J. D. (2007). Matplotlib: A 2D graphics environment. Computing in Science & Engineering, 9(3):90–95.

Hyvärinen, A., Sasaki, H., and Turner, R. (2019). Nonlinear ICA using auxiliary variables and generalized contrastive learning. In International Conference on Artificial Intelligence and Statistics, volume 22, pages 859–868.

Inan, H., Erdogdu, M. A., and Schnitzer, M. (2017). Robust estimation of neural signals in calcium imaging. In Advances in Neural Information Processing Systems, volume 30, pages 2901–2910.

Inan, H., Schmuckermair, C., Tasci, T., Ahanonu, B., Hernandez, O., Lecoq, J., Dinc, F., Wagner, M. J., Erdogdu, M., and Schnitzer, M. J. (2021). Fast and statistically robust cell extraction from large-scale neural calcium imaging datasets. bioRxiv:2021.03.24.436279.

Jones, E., Oliphant, T., Peterson, P., et al. (2001). SciPy: Open source scientific tools for Python.

Jordan, R. and Keller, G. B. (2020). Opposing influence of top-down and bottom-up input on excitatory layer 2/3 neurons in mouse primary visual cortex. Neuron.

Keller, G. B., Bonhoeffer, T., and Hübener, M. (2012). Sensorimotor mismatch signals in primary visual cortex of the behaving mouse. Neuron, 74(5):809–815.

Kerlin, A., Boaz, M., Flickinger, D., MacLennan, B. J., Dean, M. B., Davis, C., Spruston, N., and Svoboda, K. (2019). Functional clustering of dendritic activity during decision-making. Elife, 8.

Kumaran, D. and Maguire, E. A. (2006). An unexpected sequence of events: mismatch detection in the human hippocampus. PLoS Biology, 4(12):e424.

Larkum, M. E. (2013a). A cellular mechanism for cortical associations: an organizing principle for the cerebral cortex. Trends in Neurosciences, 36(3):141–151.

Larkum, M. E. (2013b). The yin and yang of cortical layer 1. Nature Neuroscience, 16(2):114–115.

Larkum, M. E., Waters, J., Sakmann, B., and Helmchen, F. (2007). Dendritic spikes in apical dendrites of neocortical layer 2/3 pyramidal neurons. The Journal of Neuroscience, 27(34):8999–9008.

Larochelle, H. and Hinton, G. E. (2010). Learning to combine foveal glimpses with a third-order Boltzmann machine. In Advances in Neural Information Processing Systems, volume 23, pages 1243–1251.

Lee, T. S. and Mumford, D. (2003). Hierarchical Bayesian inference in the visual cortex. Journal of the Optical Society of America A, 20(7):1434–1448.

Leinweber, M., Ward, D. R., Sobczak, J. M., Attinger, A., and Keller, G. B. (2017). A sensorimotor circuit in mouse cortex for visual flow predictions. Neuron, 95(6):1420–1432.

Lotter, W., Kreiman, G., and Cox, D. (2016). Deep predictive coding networks for video prediction and unsupervised learning. arXiv preprint arXiv:1605.08104.

Marques, T., Nguyen, J., Fioreze, G., and Petreanu, L. (2018). The functional organization of cortical feedback inputs to primary visual cortex. Nature Neuroscience, 21(5):757–764.

Mathis, A., Mamidanna, P., Cury, K. M., Abe, T., Murthy, V. N., Mathis, M. W., and Bethge, M. (2018). DeepLabCut: Markerless pose estimation of user-defined body parts with deep learning. Nature Neuroscience, 21(9):1281–1289.

MATLAB (2019). 9.6.0.2030181 (R2019a). The MathWorks Inc., Natick, MA.

McKinney, W. et al. (2010). Data structures for statistical computing in Python. In Proceedings of the 9th Python in Science Conference, volume 445, pages 51–56. Austin, TX.

Millman, D. J., Ocker, G. K., Caldejon, S., Larkin, J. D., Lee, E. K., Luviano, J., Nayan, C., Nguyen, T. V., North, K., Seid, S., et al. (2020). VIP interneurons in mouse primary visual cortex selectively enhance responses to weak but specific stimuli. Elife, 9:e55130.

Montijn, J. S., Meijer, G. T., Lansink, C. S., and Pennartz, C. M. (2016). Population-level neural codes are robust to single-neuron variability from a multidimensional coding perspective. Cell Reports, 16(9):2486–2498.

Murayama, M., Pérez-Garci, E., Nevian, T., Bock, T., Senn, W., and Larkum, M. E. (2009). Dendritic encoding of sensory stimuli controlled by deep cortical interneurons. Nature, 457(7233):1137.

Niell, C. M. and Stryker, M. P. (2010). Modulation of visual responses by behavioral state in mouse visual cortex. Neuron, 65(4):472–479.

Orlova, N., Tsyboulski, D., Najafi, F., Seid, S., Kivikas, S., Griffin, F., Leon, A., L’Heureux, Q., North, K., Swapp, J., et al. (2020). Multiplane mesoscope reveals distinct cortical interactions following expectation violations. bioRxiv:2020.10.06.328294.

Pedregosa, F., Varoquaux, G., Gramfort, A., Michel, V., Thirion, B., Grisel, O., Blondel, M., Prettenhofer, P., Weiss, R., Dubourg, V., Vanderplas, J., Passos, A., Cournapeau, D., Brucher, M., Perrot, M., and Duchesnay, E. (2011). Scikit-learn: Machine learning in Python. Journal of Machine Learning Research, 12:2825–2830.

Peirce, J. W. (2009). Generating stimuli for neuroscience using PsychoPy. Frontiers in Neuroinformatics, 2:10.

Poort, J., Khan, A. G., Pachitariu, M., Nemri, A., Orsolic, I., Krupic, J., Bauza, M., Sahani, M., Keller, G. B., Mrsic-Flogel, T. D., et al. (2015). Learning enhances sensory and multiple non-sensory representations in primary visual cortex. Neuron, 86(6):1478–1490.

Press, C., Kok, P., and Yon, D. (2020). The perceptual prediction paradox. Trends in Cognitive Sciences, 24(1):13–24.

Rao, R. P. and Ballard, D. H. (1999). Predictive coding in the visual cortex: a functional interpretation of some extra-classical receptive-field effects. Nature Neuroscience, 2(1):79–87.

Ruebel, O., Tritt, A., Dichter, B., Braun, T., Cain, N., Clack, N., Davidson, T. J., Dougherty, M., Fillion-Robin, J.-C., Graddis, N., et al. (2019). NWB:N 2.0: An accessible data standard for neurophysiology. bioRxiv:523035.

Rule, M. E., O’Leary, T., and Harvey, C. D. (2019). Causes and consequences of representational drift. Current Opinion in Neurobiology, 58:141–147.

Sacramento, J., Ponte Costa, R., Bengio, Y., and Senn, W. (2018). Dendritic cortical microcircuits approximate the backpropagation algorithm. In Advances in Neural Information Processing Systems, volume 31, pages 8721–8732.

Salkoff, D. B., Zagha, E., McCarthy, E., and McCormick, D. A. (2020). Movement and performance explain widespread cortical activity in a visual detection task. Cerebral Cortex, 30(1):421–437.

Smith, S. L., Smith, I. T., Branco, T., and Häusser, M. (2013). Dendritic spikes enhance stimulus selectivity in cortical neurons in vivo. Nature, 503(7474):115–120.

Spratling, M. W. (2017). A review of predictive coding algorithms. Brain and Cognition, 112:92–97.

Stringer, C., Pachitariu, M., Steinmetz, N., Reddy, C. B., Carandini, M., and Harris, K. D. (2019). Spontaneous behaviors drive multidimensional, brainwide activity. Science, 364(6437).

van den Oord, A., Li, Y., and Vinyals, O. (2018). Representation learning with contrastive predictive coding. arXiv preprint arXiv:1807.03748.

Van Rossum, G. and Drake, F. L. (2009). Python 3 Reference Manual. CreateSpace, Scotts Valley, CA.

Van Rossum, G. and Drake, F. L. J. (1995). Python Reference Manual. Centrum voor Wiskunde en Informatica Amsterdam.

Wayne, G., Hung, C.-C., Amos, D., Mirza, M., Ahuja, A., Grabska-Barwinska, A., Rae, J., Mirowski, P., Leibo, J. Z., Santoro, A., et al. (2018). Unsupervised predictive memory in a goal-directed agent. arXiv preprint arXiv:1803.10760.

Whittington, J. C. and Bogacz, R. (2017). An approximation of the error backpropagation algorithm in a predictive coding network with local Hebbian synaptic plasticity. Neural Computation, 29(5):1229–1262.

Whittington, J. C. and Bogacz, R. (2019). Theories of error back-propagation in the brain. Trends in Cognitive Sciences, 23(3):235–250.

Woloszyn, L. and Sheinberg, D. L. (2012). Effects of long-term visual experience on responses of distinct classes of single units in inferior temporal cortex. Neuron, 74(1):193–205.

Zmarz, P. and Keller, G. B. (2016). Mismatch receptive fields in mouse visual cortex. Neuron, 92(4):766–772.

Zylberberg, J., Murphy, J. T., and DeWeese, M. R. (2011). A sparse coding model with synaptically local plasticity and spiking neurons can account for the diverse shapes of V1 simple cell receptive fields. PLoS Computational Biology, 7(10):e1002250.

